# ALPAR: Automated Learning Pipeline for Antimicrobial Resistance

**DOI:** 10.1101/2025.07.08.663126

**Authors:** Alper Yurtseven, Roman Joeres, Olga V. Kalinina

## Abstract

The field of machine learning in antimicrobial resistance (AMR) research has experienced rapid growth, fueled by advancements in high-throughput genome sequencing and growing capacity of computational resources. However, the complexity and lack of standardized data preparation and bioinformatic analyses present significant challenges, especially to newcomers to the domain.

In response to these challenges, we introduce ALPAR (Automated Learning Pipeline for Antimicrobial Resistance), a comprehensive AMR data analysis tool covering the entire process from processing of raw genomic data to training machine learning models to interpretation of results. Our method relies on a reproducible pipeline that integrates widely used bioinformatics tools, presenting a simplified, automatic workflow specifically tailored for single-reference AMR analysis. Accepting genomic data in the form of FASTA files as input, ALPAR facilitates generation of machine learning-ready data tables and both training of machine learning and execution of genome-wide association studies (GWAS) experiments. Additionally, our tool offers supplementary functionalities such as phylogeny-based analysis of distribution of mutations, enhancing its utility for researchers. Our tool is accessible through the ALPAR GitHub page (https://github.com/kalininalab/ALPAR) and installable via conda (https://anaconda.org/kalininalab/ALPAR).

## Importance

While sequencing technologies are rapidly advancing, obtaining genome sequences even for many clinically relevant bacterial strains has become easily accessible for most researchers. Phenotypic resistance screens are still more labor- and cost-extensive, but large panels of this kind are starting to emerge (1, 2). The bioinformatics analysis of these data requires as a rule organization and optimization of multiple open-source software tools, which may present a challenge to researchers without profound programming experience. Hence, automated and semi-automated analysis pipelines could be valuable for a large clientele both among bioinformaticians and wet-lab scientists.

To address this need, we present ALPAR, a single reference bacterial genome analysis tool to predict antimicrobial resistances (AMR) and their determinants, as an option for processing bacterial genome data, creating machine learning (ML) ready data tables, and conducting machine learning analysis. ALPAR also offers important pre-processing steps, such as a correction for population structure using the phylogeny related parallelism score (PRPS) (3) and minimization of data leakage via creating challenging training/test data splits with DataSAIL (4). As a usage example, we processed 1,000 *Escherichia coli* genomes from public sources and ∼6,000 strains belonging to six species from the CAMDA 2024 Anti-Microbial Resistance Prediction Challenge (5).

## Introduction

Antimicrobial resistance (AMR) is a pressing global health concern, posing significant challenges to the effective treatment of bacterial infections and jeopardizing decades of progress in modern medicine (6–8). The rapid evolution and spreading of antimicrobial-resistant pathogens have necessitated innovative approaches to understand, monitor, and combat this growing threat (9, 10). Recently, machine learningbased methodologies have emerged as powerful tools in the study of AMR, capitalizing on advancements in highthroughput DNA sequencing technologies to elucidate complex relationships between genomic mutations and AMR; a well-known example of such relationship being the singlenucleotide polymorphisms (SNPs) in the *gyrA* gene which causes resistance towards ciprofloxacin. (2, 11, 12). The application of machine learning algorithms in AMR research holds promise for revolutionizing our understanding of bacterial resistance mechanisms and informing more targeted therapeutic interventions. By systematically analyzing vast datasets comprising genomic sequences and AMR phenotypic profiles, these algorithms can uncover subtle patterns and associations that may escape traditional analytical methods (13).

Traditionally, AMR prediction from genomic data relies on rule-based approaches and genome-wide association studies. Rule-based approaches involve applying predefined sets of rules derived from clinical guidelines (14, 15). These approaches rely on predefined criteria to identify patterns in genomic sequences or clinical data linked to resistance to specific antimicrobial agents. For instance, certain genetic mutations or the presence of resistance genes (e.g., *bla* genes for beta-lactam resistance (16)) can be flagged as indicators of resistance. However, this method requires extensive and detailed knowledge of specific resistance mechanisms and frequent manual updates, making it challenging to apply across the diverse organisms encountered in clinical practice (17, 18). They also often struggle to predict complex resistance patterns with multiple mechanisms involved, such as for example carbapenem resistance (18–20).

Genome-wide association studies (GWAS) enhance this analysis by scanning large collections of bacterial genomes to identify genetic variants associated with resistance to antimicrobial agents (21). GWAS compares the genomes of resistant and susceptible strains to pinpoint SNPs or other genetic markers that correlate with resistance (22). This approach enables the discovery of novel resistance mechanisms and genes, providing insights into the genetic basis of AMR (23, 24). GWAS is a powerful tool for understanding how resistance evolves and spreads, aiding in the development of new diagnostic tools and therapeutic strategies to combat resistant infections (21). Therefore, combining rule-based methods and GWAS can offer a comprehensive approach to tackling antimicrobial resistance (25–27).

Machine learning (ML) approaches, on the other hand, offer the potential to accelerate the discovery of novel resistance markers, optimize treatment regimens, and mitigate the spread of resistant pathogens (28). Unlike traditional rule-based systems, which depend on predefined rules and can be limited by their static nature, ML algorithms can dynamically adapt to new data and uncover complex patterns that are not immediately apparent (29). In contrast to GWAS, ML methods are capable of identifying non-linear effects and interactions between multiple genetic markers (30). Additionally, ML can integrate and analyze large-scale, multidimensional data from diverse sources, providing a more nuanced understanding of resistance mechanisms (31). This advanced analytical power enables the development of more accurate and personalized diagnostic tools and therapeutic strategies, significantly enhancing our ability to combat resistant infections and address the evolving challenges of antimicrobial resistance (32). ML has previously been applied in the AMR research. Traditional ML algorithms, such as support vector machines (SVMs) (33), have demonstrated strong predictive performance when applied to features such as single SNPs, gene expression profiles, and gene presence–absence (GPA) patterns in *Pseudomonas aeruginosa* (2). In addition, ensemble-based methods, particularly random forests (RF) (34), have also yielded promising results in AMR prediction tasks across various bacterial species (11).

However, despite the immense potential of ML in the AMR research, several challenges hinder its widespread adoption. A notable barrier is the extend of data preparation and bioinformatic analyses required for meaningful interpretation of genomic data in the context of antimicrobial resistance (35). Researchers, particularly those new to the field, often encounter difficulties navigating the numerous tools, formats, and protocols necessary for conducting robust AMR studies. Several tools have been offered previously to detect AMR in bacterial strains. One such example is PARGT (Prediction of Antimicrobial Resistance via Game Theory) (36), an open-source software, that identifies AMR genes in both Gram-positive and Gram-negative bacteria by training an SVM with user-input sequences. It automatically generates protein features, uses a feature selection algorithm (GTDWFE) for better prediction accuracy, and allows updates with new sequences. However, the features that are considered by the model do not include individual mutations and are related to integral properties of the protein or its segments (amino acid composition, secondary structure, PSSM bigrams, composition-transition-distribution properties). Moreover, the ML engine is limited to the SVM algorithm.

Another example is KOVER-AMR (37), a whole-genome sequencing-based antimicrobial susceptibility testing (AST) tool that employs the KOVER supervised machine learning algorithm (37) to predict phenotypic resistance. It constructs interpretable, rule-based models by identifying the presence or absence of specific *k*-mers from public genotype-phenotype databases. The tool generates two types of models: Set Covering Machines (SCM) and Classification and Regression Trees (CART). However, KOVER-AMR does not support widely used machine learning algorithms such as random forest and gradient boosting trees.

To address these limitations and facilitate accessibility to ML-based AMR analysis, we present ALPAR (Automated Learning Pipeline for Antimicrobial Resistance), a comprehensive and user-friendly pipeline for conducting single-reference computational AMR analysis experiments. As genetic markers used as features for the ML methods, ALPAR employs SNPs, short insertions and deletions (indel), gene presence or absence (GPA). SNPs and indels offer high-resolution insights into genomic variation by capturing single or few base-pair differences between isolates. In contrast, GPA features summarize broader functional variation, reducing dimensionality and enhancing interpretability, though they sacrifice fine-scale resolution and overlook non-coding regions (38). ALPAR uses strength of both approaches by combining them. Our method establishes a standardized framework for identifying genetic variants such as SNPs and indels, while also enabling the detection of GPA across multiple samples. Our approach is grounded in the integration of commonly used bioinformatic tools into a single pipeline. By streamlining the workflow and providing clear guidelines, our pipeline aims to empower researchers to embark on AMR research endeavors with confidence and efficiency.

## Results

### A. Design and implementation

ALPAR (Automated Learning Pipeline for Antimicrobial Resistance) is a software package implemented in Python for easy readability and extendability. It links several open-source bioinformatics software tools to allow users to conduct single-reference AMR experiments in one workflow.

As input, ALPAR accepts genomic fasta files and reference genome In order to create binary mutation tables, ALPAR employs the following pipeline: Snippy (39) is used to call variants for each strain against a reference genome, Prokka (40) annotates each genome using a user-defined protein database, CD-HIT clusters annotated genomes to determine GPA information with minimal resource requirements, and Panaroo (41) provides more detailed GPA information. For building the phylogenetic tree of all genomes, ALPAR offers two options: MashTree (42) for an alignment-free phylogenetic tree, and the PanACoTA pipeline (43) for building an alignment-based phylogenetic tree. As intermediate steps, ALPAR allows to calculate PRPS (phylogeny-related parallelism scores) (3) that allows to prioritize genetic markers that are weakly correlated with the phylogeny (as they are assumed to more likely be functionally relevant), and compute splits for further training ML models with DataSAIL (4) to mitigate the effects of information leakage. Then several ML models are trained: RF, SVM and gradient boosting (GB) depending on user’s configuration. While training the ML models, ALPAR employs a grid search method for easy parameter optimization to find the best models.

#### Use case: *Escherichia coli* AMR data from PATRIC Database (1)

To showcase our approach, we applied the ALPAR pipeline to publicly available *E. coli* dataset of 1000 genomes from the PATRIC database (1). We selected *E. coli* as our example organism because it is a well-established bacterial model (44). The genomes from the dataset were already annotated as resistant or susceptible; therefore, we did not need to perform any pre-preparation steps. We have used the automatix option to run the entire pipeline with the following command:

~~~
alpar automatix -i Escherichia/ -o
Escherichia_out/ --reference
Escherichia_Reference.gbff --
custom_database
Escherichia_Proteins.fasta --
threads 16 --ram 128 --
ml_algortihm rf --fast
~~~

Automatix is an automated function to run ALPAR. The `-i` option specifies the input path, which can be either a folder or a text file where each line contains the full path of a strain that the user wants to process. The `-o` option specifies the output folder path. The `--reference` option specifies the reference genome GBFF file. The `--custom_database` option is not required but is strongly recommended to achieve more accurate GPA information; it should be FASTA file of all the proteins that exist in the given species (these can be collected from UniProt (45)). The `--threads` option allows the user to specify the number of threads to be used for the run, and the `--ram` option limits memory usage (value given in gigabytes). The `--ml_algorithm` option specifies which machine learning algorithm the user wants to run; the default option is to run random forest, gradient boosting, and support vector machine algorithms. The `--fast` option skips the alignment-based phylogenetic tree and generates it using an alignment-free method. Full documentation can be found on GitHub (https://github.com/kalininalab/ALPAR) and Read the Docs page (https://alpar.readthedocs.io/en/latest/).

Running the full pipeline took ∼35 hours wall time on a 2x AMD EPYC 7601 32-Core Processors (128 threads total) and 1TiB of RAM, running Linux (CentOS 8, kernel 4.18.0). 56 strains out of 1,000 were skipped because the variant calling step failed. A later manual check showed that the genomes of these strains were either 95% shorter or 10% longer than the reference genome *Escherichia coli* str. K-12 substr. MG1655 (accession number NC_000913 (46)). 944 strains that had been screened against ciprofloxacin were used in later steps. The pipeline has generated 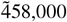 features. As a result of the automatix pipeline, our random forest classification model achieved an MCC of 0.891. Feature importance analysis, conducted by means of the permutation feature importance routine, yielded two significant mutations: the S80I mutation in the ParC protein (DNA topoisomerase 4 subunit A, Figure 2), a missense mutation with an importance value 0.153, and the S83L mutation in the GyrA protein (DNA gyrase subunit A, Figure 2), a missense mutation with an importance value 0.115. Both mutations are well-known resistance determinants towards fluoroquinolone drugs, of which ciprofloxacin is an example (47).

**Fig. 1.**
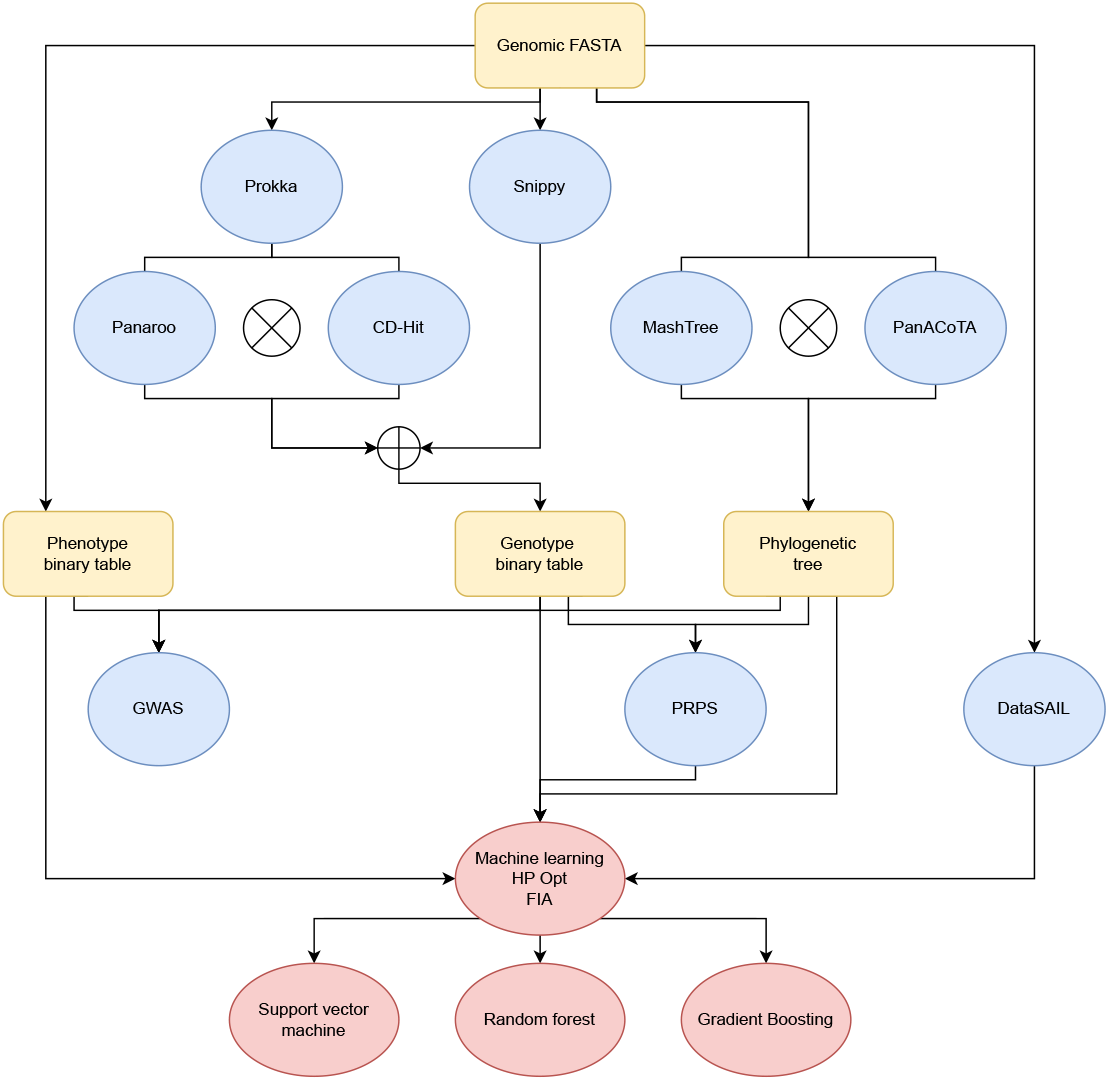
General overview of the ALPAR’s pipeline

**Fig. 2.**
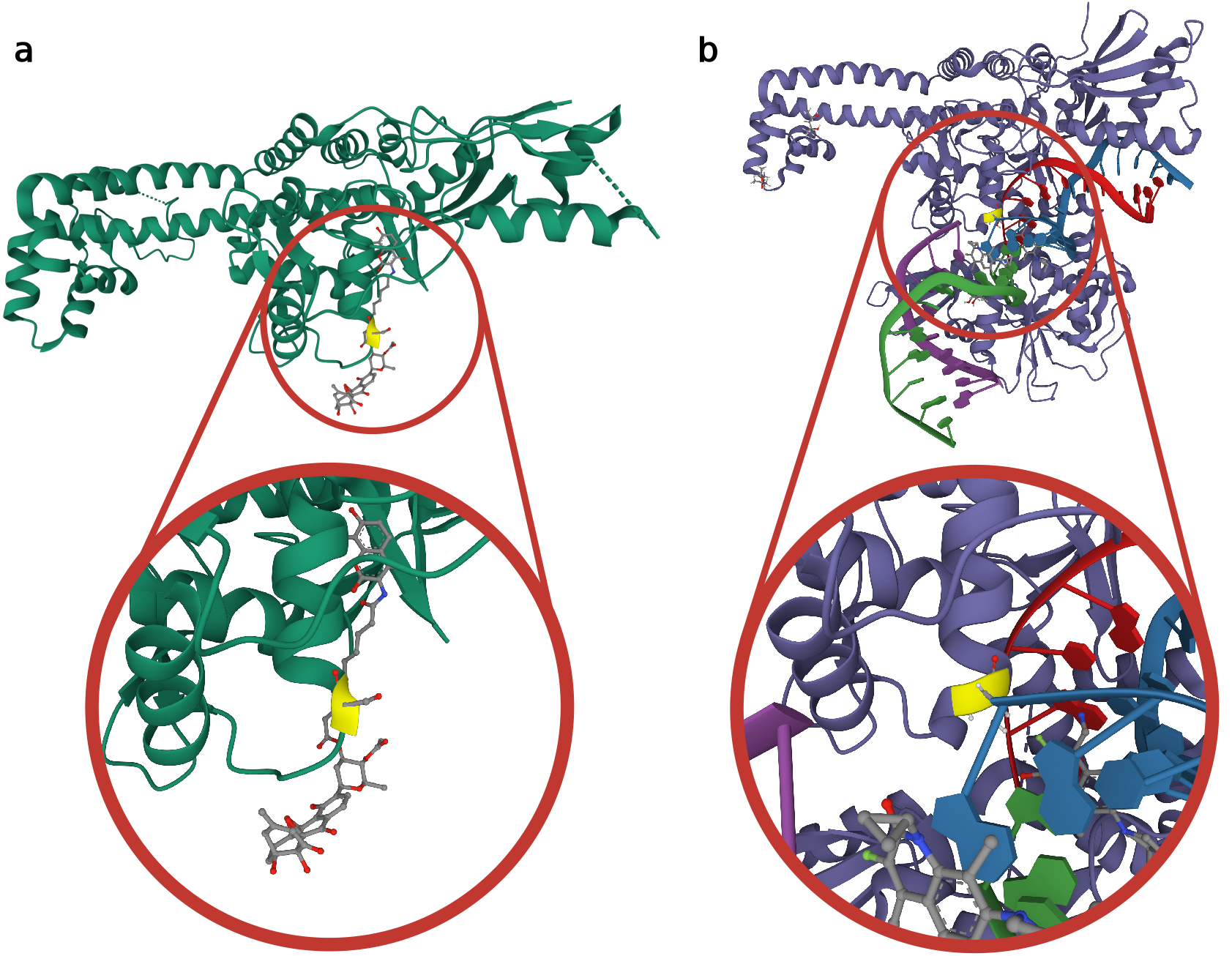
Mapping of mutations with the highest weight in the feature importance analysis for the *E. coli* dataset. **(a)** The S83L mutation in GyrA; mapping based on the experimentally resolved structure of the *E. coli* GyrA, PDB ID: 4CKL. **(b)** The S80I mutation in ParC; mapping based on the experimentally resolved structure of the *P. putida* ParC, PDB ID: 6BQ9. Mutated amino acids are shown in yellow and visualized using StructMAn Web (49).

To further investigate the mechanism of resistance, we analyzed the identified mutations with StructMAn 2.0 Web, a web-based tool that annotates non-synonymous singlenucleotide variants (nsSNVs) by analyzing their structural context within 3D protein structures, including interactions with other macromolecules and ligands (48) (49). Both positions are structurally homologous and were classified as interacting with a ligand, which explains the resistance: the change of a serine residues to a hydrophobic disrupts the interaction with the drug (Figure 2).

#### Use case: CAMDA Anti-Microbial Resistance Prediction Challenge – 2024

In the 2024 CAMDA challenge (5), we focused on building ML models to predict the AMR status of 1,820 bacterial strains that belong to seven different species (*Campylobacter jejuni, Campylobacter coli, Escherichia coli, Klebsiella pneumoniae, Neisseria gonorrhoeae, Pseudomonas aeruginosa, Salmonella enterica*). For these strains, the AMR status towards two drugs (meropenem, ciprofloxacin) was provided by the challenge organizers.

Here we also employed the automatix module of ALPAR utilizing the random forest method. For each bacterial–antibiotic pair in the dataset, two models were trained: one including all the features and one only using the top 30% of the features ranked according to their PRPS score that penalizes variants that are localized to a single branch on the strains’ phylogenetic tree. Predictions on the test set were generated using both models and yielded identical resistance status classifications across all 1,820 test strains. The prediction results were submitted to the CAMDA challenge, achieving a score of 83 out of 100, which was the winning score in the challenge.

#### Use case: Large-scale AMR screening across 11 species from the PATRIC database

To further demonstrate the scalability and generalizability of ALPAR, we extended our evaluation to a large-scale antimicrobial resistance screening across multiple species using data from the PATRIC database (1). In this study, we curated genome and resistance phenotype data for 11 bacterial species, screened against 37 unique antibiotics, resulting in 79 distinct bacteria–antibiotic pairs.

Each dataset was processed using ALPAR’s automatix module with the same parameters as in the previous use cases. For each pair, machine learning models were trained to predict resistance status of strains by using SNPs, indels & GPA features. Model performance metrics, F1-score and Matthews correlation coefficient (MCC), were recorded, and gini importance based feature importance analyses were conducted to identify the potential causative genetic markers.

Across the 79 bacteria–antibiotic pairs, our models achieved strong predictive performance, with several models reaching MCC values above 0.85 (Figure 3, detailed scores and results of the feature importance analysis can be found in Supplementary Table 1).

**Fig. 3.**
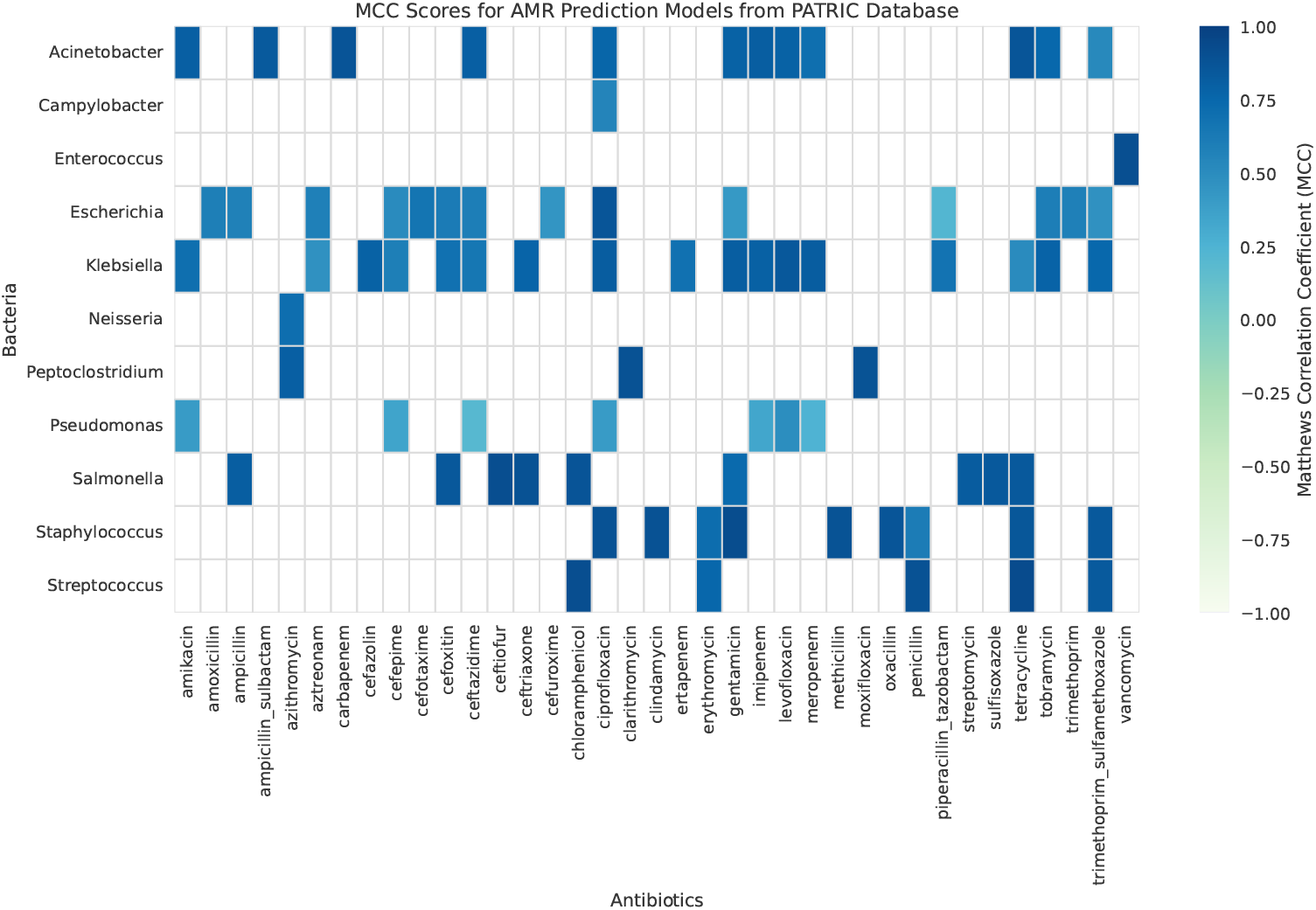
Heatmap of MCC scores for AMR prediction models across different bacteria-antibiotic combinations from PATRIC database (1). Color intensity indicates model performance, with darker blue representing better predictive accuracy and light green representing poor performance.

## Discussion

ALPAR is a flexible tool for antimicrobial-resistant bacterial genomes. It can be run on a laptop for a small number of bacterial samples, but, critically, the underlying multi-threading allows it to make efficient use of large clusters and cloud-computing infrastructures to process the many thousands of genomes. It can be run with any bacterium and antibiotic resistance profile as long as the genomic sequence and reference genome are provided. For users who are not familiar with antimicrobial-resistant bacterial genomic tools or machine learning, and/or who require a standardized pipeline, ALPAR is a one-stop tool that can be easily deployed using conda. For researchers with experience in the field, ALPAR also offers analysis with other tools recently developed, such as the PRPS score and DataSAIL, to improve machine learning models’ prediction performances by reducing the effects of phylogenetic structures and information leakage on the models to address issues mentioned in recent paper by James et al. (2025) (38).

One potential limitation of ALPAR is its reliance on a predefined reference strain. In real-world applications it can be difficult to single out one particular strain as a reference, and multi-reference graph-based tools have emerged (50). The use of a single reference genome ensures consistency across all strains and offers an easy solution to genome sequencing quality issues. Single-reference variant calling and GPA analysis provide an effective approach for genomic comparisons by aligning all samples to a single reference genome. This approach has been applied and tested in several studies on AMR predictions. For example, Khaledi et al. (2) used single-reference genome variant calling for *P. aeruginosa* strains screened against four different antibiotics, using the *P. aeruginosa* PA14 reference strain. Similarly, Ren et al. (11) applied the single-reference approach to *E. coli* strains screened against four antibiotics, using the *E. coli* K-12 reference strain. Nevertheless, extending ALPAR with a multireference variant caller is also a conceivable future direction of research.

ML methods have firmly established themselves in the AMR research, although they have some inherent limitations, which mainly stem from complex bacterial genomics (38).Through our contributions, we provide an easy-to-use comprehensive pipeline for creating versatile ML models for predicting AMR from genomic data and thus seek to progress in the ongoing battle against antimicrobial resistance.

## Materials and Methods

### The ALPAR package

ALPAR is an open-source bioinformatics software package that accepts (i) genomic data in FASTA-formatted files of assembled bacterial whole genome sequences and (ii) a reference genome to analyze. In return, it creates machine learning ready mutation and GPA binary tables (see Fig. 1). These feature tables are then used to train predictive models. In addition to the trained models, ALPAR also outputs model performance metrics and feature importance analysis, helping users interpret the biological relevance of genomic variations. In the following sections, we present the individual steps of the pipeline.

### A.1. Creating binary mutation and GPA tables

Typically, for AMR analysis, genomic sequences are transformed into binary mutation tables and tables that report GPA information. Binary mutation tables summarize information on genomic variation by converting SNPs, indels, and complex genomic rearrangements into binary features: presence or absence of the variant compared to the reference strain. In ALPAR, we focus on SNPs and indels. GPA analysis complements this by examining the presence or absence of entire genes within genomes of individual strains compared to the reference.

ALPAR creates these tables in a three-step process. (i) Snippy (39) is used to call variants from strains against a reference genome, then (ii) Prokka (40) annotates genomes with an optional user-defined genus protein database to ascertain GPA information. Lastly, (iii) CD-HIT (51) or Panaroo (41) cluster these annotated genomes to extract GPA information. Additionally, ALPAR offers generation of a binary phenotype table from all these steps or only variants automatically.

### A.2. Phylogeny pipeline

Phylogenetic information can be useful for antimicrobial resistance (AMR) analysis: due to the typically low recombination rates in bacteria (52), many genetic markers may appear to be associated with AMR as a consequence of linkage disequilibrium rather than direct causal relationships (38). ALPAR offers both alignment-free and alignment-based approaches for constructing the phylogenetic tree, allowing users to choose the method that best suits their data and analysis goals. In the alignment-free approach, ALPAR computes Mash distances (53) and constructs a phylogenetic tree using Mashtree (42). The resulting tree can be converted into a distance matrix for integration into the machine learning framework. In the alignment-based approach, ALPAR utilizes the PanACoTA pipeline (43), which includes genome quality control, pan-genome analysis, and phylogenetic tree construction based on multiple sequence alignment using IQ-TREE (54).

### A.3. Phylogeny-related parallelism score (PRPS)

Phylogenetic relations are often seen as a challenge in machine learning studies of AMR because they can make models focus on lineage-specific instead of resistance-related variants (38). To reduce this effect in machine learning analysis, ALPAR employs a method called the phylogeny-related parallelism score (PRPS) (3). PRPS ranks mutations according to their tendency to cluster within single or a few branches of the phylogenetic tree as opposed to being scattered throughout the whole tree. The latter kind of mutations, when correlated with AMR, are more likely to be causative. ALPAR can drop a user-defined percentage of low-PRPS features from the data frame to reduce the number of features considered by ML models while retaining potentially causative variants. Our suggested value is 70%, which is used as the default.

### A.4. Genome-wide association studies (GWAS) analysis

A state-of-the-art approach for identifying significant mutations associated with resistance is the genome-wide association studies (GWAS). The strong linkage disequilibrium prevalent in bacterial genomes led to establishing a specialized framework for bacterial GWAS (55). To find strong associations between genotype and phenotype among bacterial strains, ALPAR leverages GWAS analysis with Pyseer (56) that employs linear mixed models to reduce the strong linkage disequilibrium effect of bacteria.

### A.5. Machine learning

Machine learning techniques have emerged as essential tools in the study of antimicrobial resistance (AMR), offering versatile advantages across various aspects of research (57). Even though GWAS can achieve this to some extent, machine learning models offer an improved approach to understanding multi-locus genetic variants and their interactions in predicting complex traits (58). Although machine learning tools are easily accessible, there are important steps that should be followed when applying them to bacterial data due to their characteristics, such as strong linkage disequilibrium and population structure. That makes it challenging for new scientists in the field. ALPAR implements an easy-to-use command line interface for commonly used machine learning classification algorithms: Support vector machine (SVM), random forest (RF) and gradient boosting (GB), employing grid search for hyperparameter optimization.

In addition, ALPAR addresses the critical issue of information leakage in machine learning, which can lead to inflated performance reports. Information leakage occurs when information from the test data inadvertently influences model training, diminishing performance when applied in a realworld scenario. To mitigate this risk, ALPAR employs DataSAIL (4), which strategically splits data into training, validation, and test sets to maximize the distance between the sets. We use genome distance matrices created by Mash (53) to this end. By incorporating DataSAIL, ALPAR enhances the reliability and robustness of its predictive models, thereby facilitating accurate classification of antibiotic resistance patterns and identification of crucial genomic features.

The latter is achieved by implementing feature importance analysis to detect features that most influence the AMR status of a bacterium. ALPAR employs both permutation-based feature importance analysis and Gini importance (34, 59).

### A.6. Prediction

Predicting AMR status of new strains represents the key application of a trained machine learning model in the AMR research. ALPAR offers users to predict the AMR status of new strains by utilizing pre-generated models. The prediction submodule accepts new genomes in the form of genomic FASTA files and employs the same pipeline for creating the binary variant and GPA tables. Subsequently, the submodule utilizes the saved models to predict the AMR outcomes for these new strains.

## ACKNOWLEDGEMENTS

We would like to thank Dr. Alexander Gress for stimulating discussions and in particular for naming the tool. Amay Ajaykumar Agrawal for discussion. A.Y. has been partially funded by the HelmholtzAI project AMR-XAI and the BMBF project Sys_CARE (project ID 01ZX1908C). R.J. has been partially funded by the HelmholtzAI project XAI-Graph. O.V.K. acknowledges funding from the Klaus Faber Foundation.

## Supplementary Note 1: Data Availability Statement

Genomic files for *E. coli* samples used in this study were obtained from the PATRIC Database (1). The code and software are available at GitHub (https://github.com/kalininalab/ALPAR) and conda (https://anaconda.org/kalininalab/alpar).

