## Supplementary Table 1 for "ALPAR: Automated Learning Pipeline for Antimicrobial Resistance"

### Supplementary 1. PATRIC Database Machine Learning & FIA Results

#### Campylobacter jejuni - ciprofloxacin

| Bacteria | Antibiotic | F1 Score | MCC |
|---|---|---|---|
| Campylobacter jejuni | ciprofloxacin | 0.667 | 0.553 |

Table 1: Feature Importance for Campylobacter jejuni - ciprofloxacin

| Feature Name | Gini Importance | Mutation Annotation | Gene |
|---|---|---|---|
| '959966','G:A','snp' | 0.007 | missense_variant c.257C>T p.Thr86Ile | gyrA |
| '1239397','A:G','snp' | 0.004 | synonymous_variant c.255T>C p.Ser85Ser | |
| '140005','A:G','snp' | 0.003 | synonymous_variant c.186A>G p.Gly62Gly | |
| '1451635','C:T','snp' | 0.002 | synonymous_variant c.36C>T p.Leu12Leu | |
| '249956','T:C','snp' | 0.002 | synonymous_variant c.552T>C p.Phe184Phe | |
| '1223488','G:A','snp' | 0.002 | synonymous_variant c.147C>T p.Ser49Ser | accC |
| '72331','CCA:TAG','complex' | 0.002 | missense_variant c.327_329delCCAinsTAG p.His110Ser | |
| '1239505','C:T','snp' | 0.002 | synonymous_variant c.147G>A p.Val49Val | |
| '985487','G:A','snp' | 0.002 | synonymous_variant c.1524C>T p.Tyr508Tyr | cjeI |
| '1608923','T:C','snp' | 0.002 | synonymous_variant c.540A>G p.Glu180Glu | topA |

#### Enterococcus faecalis - vancomycin

| Bacteria | Antibiotic | F1 Score | MCC |
|---|---|---|---|
| Enterococcus faecalis | vancomycin | 0.963 | 0.900 |

Table 2: Feature Importance for Enterococcus faecalis - vancomycin

| Feature Name | Gini Importance | Mutation Annotation | Gene |
|---|---|---|---|
| OI6YUCZY_POIILDGO_02470_recombinase_family_protein | 0.021 | | |

**Table 2 – continued from previous page**

| Feature Name | Gini Importance | Mutation Annotation | Gene |
|---|---|---|---|
| OI6YUCZY_POIILDGO_02471_hypothetical_protein | 0.019 | | |
| OI6YUCZY_POIILDGO_02822_D_alanine__D_alanine_ligase | 0.017 | | |
| OI6YUCZY_POIILDGO_02466_IS21_family_transposase_ISPpu7 | 0.014 | | |
| OI6YUCZY_POIILDGO_02464_hypothetical_protein | 0.014 | | |
| 3X4O2J89_FFOPJHCL_02578_Cadmium_resistance_transcriptional_regulatory_protein_CadC | 0.013 | | |
| OI6YUCZY_POIILDGO_02469_Histidine_biosynthesis_bifunctional_protein_HisB | 0.012 | | |
| O5FE2XZG_EGNPMKAE_00545_hypothetical_protein | 0.011 | | |
| 17GY39TG_AHHBCFLI_02680_Adaptive_response_sensory_kinase_SasA | 0.011 | | |
| OI6YUCZY_POIILDGO_02124_hypothetical_protein | 0.011 | | |

#### Escherichia coli - ciprofloxacin

| Bacteria | Antibiotic | F1 Score | MCC |
|---|---|---|---|
| Escherichia coli | ciprofloxacin | 0.898 | 0.861 |

Table 3: Feature Importance for Escherichia coli - ciprofloxacin

| Feature Name | Gini Importance | Mutation Annotation | Gene |
|---|---|---|---|
| '2339173','G:A','snp' | 0.016 | missense_variant c.248C>T p.Ser83Leu | gyrA |
| '2028695','GATA:AATG','complex' | 0.010 | synonymous_variant c.669_672delTATCinsCATT p.225 | yedI |

**Table 3 – continued from previous page**

| Feature Name | Gini Importance | Mutation Annotation | Gene |
|---|---|---|---|
| '3140341','C:T','snp' | 0.009 | synonymous_variant c.459G>A p.Leu153Leu | hybE |
| '1512011','A:G','snp' | 0.009 | missense_variant c.358A>G p.Thr120Ala | ydcS |
| '4477436','T:C','snp' | 0.008 | synonymous_variant c.876A>G p.Glu292Glu | argI |
| '3543049','T:C','snp' | 0.008 | | |
| '3143619','GGTC:AGTT','complex' | 0.008 | synonymous_variant c.543_546delGACCinsAACT p.183 | hybB |
| '3599473','AATGCCG:GATACCA','complex' | 0.006 | synonymous_variant c.180_186delCGGCATTinsTGGTATC p.63 | livJ |
| '4460251','T:G','snp' | 0.005 | synonymous_variant c.114A>C p.Thr38Thr | nrdG |
| '249270','A:G','snp' | 0.005 | | |

#### Escherichia coli - cefoxitin

| Bacteria | Antibiotic | F1 Score | MCC |
|---|---|---|---|
| Escherichia coli | cefoxitin | 0.682 | 0.611 |

Table 4: Feature Importance for Escherichia coli - cefoxitin

| Feature Name | Gini Importance | Mutation Annotation | Gene |
|---|---|---|---|
| '1570669','T:C','snp' | 0.006 | synonymous_variant c.1377A>G p.Gln459Gln | gadB |
| '2006761','C:T','snp' | 0.004 | synonymous_variant c.606C>T p.Gly202Gly | amyA |
| '3543049','T:C','snp' | 0.004 | | |
| '576547','G:A','snp' | 0.004 | synonymous_variant c.279C>T p.Ser93Ser | nmpC |
| '3302052','ATA:GTG','complex' | 0.004 | missense_variant c.568_570delATAinsGTG p.Ile190Val | ubiU |
| '1378744','T:C','snp' | 0.004 | synonymous_variant c.861T>C p.Asn287Asn | ycjT |
| '1759198','G:A','snp' | 0.004 | synonymous_variant c.93C>T p.Gly31Gly | sufE |
| R9Z6SGR2_HGDCBKLK_04032_hypothetical_protein | 0.004 | | |
| '4437213','G:T','snp' | 0.004 | synonymous_variant c.459G>T p.Val153Val | cysQ |
| '3305916','A:T','snp' | 0.003 | | |

#### Escherichia coli - tobramycin

| Bacteria | Antibiotic | F1 Score | MCC |
|---|---|---|---|
| Escherichia coli | tobramycin | 0.679 | 0.596 |

Table 5: Feature Importance for Escherichia coli - tobramycin

| Feature Name | Gini Importance | Mutation Annotation | Gene |
|---|---|---|---|
| ’174241’,’C:T’,’snp’ | 0.007 | synonymous_variant c.642G>A p.Pro214Pro | hemL |
| ’1512011’,’A:G’,’snp’ | 0.007 | missense_variant c.358A>G p.Thr120Ala | ydcS |
| ’348879’,’T:C’,’snp’ | 0.006 | synonymous_variant c.198T>C p.Asp66Asp | prpB |
| 4W6EIH06_ EJGMIGKJ_04445_ Aminoglycoside_N_6_ __acetyltransferase_ type_1 | 0.006 | | |
| ’2339173’,’G:A’,’snp’ | 0.006 | missense_variant c.248C>T p.Ser83Leu | gyrA |
| ’576547’,’G:A’,’snp’ | 0.006 | synonymous_variant c.279C>T p.Ser93Ser | nmpC |
| ’1956927’,’G:T’,’snp’ | 0.005 | synonymous_variant c.81C>A p.Leu27Leu | torZ |
| ’3583211’,’G:T’,’snp’ | 0.005 | missense_variant c.174G>T p.Lys58Asn | yrhA |
| ’1584606’,’G:A’,’snp’ | 0.005 | synonymous_variant c.1881C>T p.Ser627Ser | ydeP |
| ’3232444’,’T:C’,’snp’ | 0.005 | synonymous_variant c.780T>C p.Ile260Ile | fadH |

#### Escherichia coli - ceftazidime

| Bacteria | Antibiotic | F1 Score | MCC |
|---|---|---|---|
| Escherichia coli | ceftazidime | 0.655 | 0.600 |

Table 6: Feature Importance for Escherichia coli - ceftazidime

| Feature Name | Gini Importance | Mutation Annotation | Gene |
|---|---|---|---|
| ’3267775’,’G:A’,’snp’ | 0.008 | | |
| ’2824803’,’G:A’,’snp’ | 0.007 | synonymous_variant c.774C>T p.Asn258Asn | mltB |
| ’747996’,’AAG: TAA’,’complex’ | 0.007 | synonymous_variant c.985_987delCTTinsTTA p.330 | ybgO |
| ’840713’,’G:A’,’snp’ | 0.005 | synonymous_variant c.819C>T p.Gly273Gly | fiu |
| ’3877647’,’AGATTT: GGAGTC’,’complex’ | 0.005 | | |

| Feature Name | Gini Importance | Mutation Annotation | Gene |
|---|---|---|---|
| VIOXH024_NLIKFJMI_04760_IS1_family_protein_InsB | 0.005 | | |
| '3591390','C:T','snp' | 0.005 | synonymous_variant c.936G>A p.Val312Val | ugpB |
| '914594','C:T','snp' | 0.005 | synonymous_variant c.264G>A p.Pro88Pro | lysO |
| '3230024','T:C','snp' | 0.005 | synonymous_variant c.1137T>C p.Asp379Asp | ygjK |
| '3070780','G:A','snp' | 0.005 | synonymous_variant c.465C>T p.Ile155Ile | fbaA |

#### Escherichia coli - aztreonam

| Bacteria | Antibiotic | F1 Score | MCC |
|---|---|---|---|
| Escherichia coli | aztreonam | 0.706 | 0.584 |

Table 7: Feature Importance for Escherichia coli - aztreonam

| Feature Name | Gini Importance | Mutation Annotation | Gene |
|---|---|---|---|
| '1749350','G:A','snp' | 0.008 | synonymous_variant c.210C>T p.Ser70Ser | ydhT |
| '1345540','T:C','snp' | 0.008 | synonymous_variant c.1203A>G p.Gln401Gln | pdeR |
| '172817','T:C','snp' | 0.008 | synonymous_variant c.1356T>C p.Pro452Pro | fhuB |
| '4430745','TGCCGCA:GGCTGCC','complex' | 0.008 | synonymous_variant c.882_888delTGCCGCAinsGGCTGCC p.297 | cycA |
| '4315191','C:G','snp' | 0.007 | synonymous_variant c.348G>C p.Ser116Ser | phnO |
| '1766652','C:A','snp' | 0.007 | synonymous_variant c.2034G>T p.Ala678Ala | ydiJ |
| '576547','G:A','snp' | 0.007 | synonymous_variant c.279C>T p.Ser93Ser | nmpC |
| '3981654','CACC:TACT','complex' | 0.007 | synonymous_variant c.768_771delCACCinsTACT p.258 | yifK |
| '1607713','A:T','snp' | 0.005 | missense_variant c.368A>T p.Gln123Leu | tam |
| 016JET8K_BMHFLKKD_05009_hypothetical_protein | 0.005 | | |

#### Escherichia coli - cefepime

| Bacteria | Antibiotic | F1 Score | MCC |
|---|---|---|---|
| Escherichia coli | cefepime | 0.604 | 0.506 |

Table 8: Feature Importance for Escherichia coli - cefepime

| Feature Name | Gini Importance | Mutation Annotation | Gene |
|---|---|---|---|
| '3099883','G:A','snp' | 0.008 | synonymous_variant c.846C>T p.Ala282Ala | ansB |
| '1698308','TGTC: CGTT','complex' | 0.006 | synonymous_variant c.870_873delGACAinsAACG p.292 | malI |
| '3672378','T:C','snp' | 0.006 | | |
| '576920','G:A','snp' | 0.006 | | |
| '576778','T:C','snp' | 0.005 | synonymous_variant c.48A>G p.Leu16Leu | nmpC |
| '1056352','T:A','snp' | 0.005 | missense_variant c.92T>A p.Phe31Tyr | torT |
| '3977761','T:A','snp' | 0.005 | synonymous_variant c.237T>A p.Arg79Arg | wecF |
| '4432669','T:C','snp' | 0.005 | missense_variant c.388A>G p.Ile130Val | ytfF |
| '4481237','C:T','snp' | 0.004 | synonymous_variant c.2601G>A p.Thr867Thr | valS |
| DQFCZQ1D_FAOEBIJP_04964_hypothetical_protein | 0.004 | | |

#### Escherichia coli - amoxicillin

| Bacteria | Antibiotic | F1 Score | MCC |
|---|---|---|---|
| Escherichia coli | amoxicillin | 0.813 | 0.589 |

Table 9: Feature Importance for Escherichia coli - amoxicillin

| Feature Name | Gini Importance | Mutation Annotation | Gene |
|---|---|---|---|
| BZZ8XNVZ_OAKLIGDF_05308_hypothetical_protein | 0.009 | | |
| JS39IWW3_KLOOMICH_04185_hypothetical_protein | 0.005 | | |
| OS72JI7N_GABGHPOD_04899_dihydropteroate_synthase | 0.004 | | |
| 0096HECL_ONPLLBGJ_04660_Beta_lactamase_TEM | 0.004 | | |

**Table 9 – continued from previous page**

| Feature Name | Gini Importance | Mutation Annotation | Gene |
|---|---|---|---|
| 1IS7IIN9_GKMAKHGB_04163_hypothetical_protein | 0.004 | | |
| AWKE3LSI_PLPDHGMK_04566_Tn3_family_transposase_Tn2 | 0.003 | | |
| 1068LF20_BOEPEKIJ_01628_hypothetical_protein | 0.003 | | |
| 0096HECL_ONPLLBGJ_04539_dihydropteroate_synthase | 0.003 | | |
| '3091205','C:T','snp' | 0.003 | synonymous_variant c.72C>T p.Ala24Ala | rsmE |
| AZ3YBWD7_DDHODMAP_04466_Regulatory_protein_RepA | 0.003 | | |

#### Escherichia coli - trimethoprim

| Bacteria | Antibiotic | F1 Score | MCC |
|---|---|---|---|
| Escherichia coli | trimethoprim | 0.733 | 0.582 |

Table 10: Feature Importance for Escherichia coli - trimethoprim

| Feature Name | Gini Importance | Mutation Annotation | Gene |
|---|---|---|---|
| 0096HECL_ONPLLBGJ_04539_dihydropteroate_synthase | 0.006 | | |
| '3092333','G:A','snp' | 0.005 | synonymous_variant c.456G>A p.Glu152Glu | gshB |
| '1552419','TAA:CAG','complex' | 0.005 | synonymous_variant c.262_264delTTAinsCTG p.89 | yddM |
| 0096HECL_ONPLLBGJ_04509_Antitoxin_PemI | 0.004 | | |
| '3096249','C:T','snp' | 0.004 | synonymous_variant c.430C>T p.Leu144Leu | yggT |
| 0096HECL_ONPLLBGJ_04611_Aminoglycoside_3__phosphotransferase | 0.004 | | |
| JR3PQ66I_FPIPAIKB_04571_Streptomycin_3__adenylyltransferase | 0.004 | | |
| '3044438','CGCT:TGCC','complex' | 0.004 | synonymous_variant c.777_780delCGCTinsTGCC p.261 | bglA |
| '725612','A:G','snp' | 0.003 | synonymous_variant c.1425T>C p.Arg475Arg | kdpB |

**Table 10 – continued from previous page**

| Feature Name | Gini Importance | Mutation Annotation | Gene |
|---|---|---|---|
| '3049115','AT: A','del' | 0.003 | | |

#### Escherichia coli - piperacillin_tazobactam

| Bacteria | Antibiotic | F1 Score | MCC |
|---|---|---|---|
| Escherichia coli | piperacillin_tazobactam | 0.286 | 0.224 |

Table 11: Feature Importance for Escherichia coli - piperacillin_tazobactam

| Feature Name | Gini Importance | Mutation Annotation | Gene |
|---|---|---|---|
| 0OPRY3ZM_PKJIMEEA_04269_outer_membrane_porin_C | 0.002 | | |
| 0OPRY3ZM_PKJIMEEA_04469_IS1_family_protein_InsA | 0.002 | | |
| '3278576','C:T','snp' | 0.002 | synonymous_variant c.90G>A p.Gly30Gly | agaR |
| 1INNSZVT_PNLHLPDI_05276_IS21_family_transposase_ISKpn7 | 0.002 | | |
| '1928192','G:A','snp' | 0.002 | synonymous_variant c.648C>T p.Ile216Ile | ptrB |
| '3142204','G:A','snp' | 0.002 | synonymous_variant c.786C>T p.Thr262Thr | hybC |
| '1478750','A:G','snp' | 0.002 | synonymous_variant c.525A>G p.Ala175Ala | ynbB |
| '3276367','C:T','snp' | 0.001 | synonymous_variant c.1086C>T p.Asp362Asp | garD |
| '3873439','G:A','snp' | 0.001 | synonymous_variant c.174C>T p.Asp58Asp | dgoA |
| '574184','GTC: TGT','complex' | 0.001 | missense_variant c.229_231delGTCinsTGT p.Val77Cys | quuD |

#### Escherichia coli - gentamicin

| Bacteria | Antibiotic | F1 Score | MCC |
|---|---|---|---|
| Escherichia coli | gentamicin | 0.491 | 0.416 |

Table 12: Feature Importance for Escherichia coli - gentamicin

| Feature Name | Gini Importance | Mutation Annotation | Gene |
|---|---|---|---|
| `03L1GF6O_LBBKHAIB_04655_SPbeta_prophage_derived_aminoglycoside_N_3___acetyltransferase_like_protein_YokD` | 0.011 | | |
| `'76262','G:T','snp'` | 0.007 | synonymous_variant c.1038C>A p.Arg346Arg | sgrR |
| `'1148839','C:T','snp'` | 0.007 | synonymous_variant c.81C>T p.Asp27Asp | fabH |
| `'3176185','G:A','snp'` | 0.006 | synonymous_variant c.649C>T p.Leu217Leu | cpdA |
| `'2339166','G:A','snp'` | 0.005 | synonymous_variant c.255C>T p.Val85Val | gyrA |
| `'3390161','A:G','snp'` | 0.005 | synonymous_variant c.642A>G p.Pro214Pro | aaeR |
| `'3877647','AGATTT:GGAGTC','complex'` | 0.004 | | |
| `'3134487','A:G','snp'` | 0.004 | synonymous_variant c.357A>G p.Gln119Gln | yghT |
| `'310615','C:T','snp'` | 0.004 | synonymous_variant c.57G>A p.Val19Val | ecpA |
| `'817184','T:G','snp'` | 0.004 | synonymous_variant c.141T>G p.Leu47Leu | moaA |

#### Escherichia coli - cefotaxime

| Bacteria | Antibiotic | F1 Score | MCC |
|---|---|---|---|
| Escherichia coli | cefotaxime | 0.703 | 0.657 |

Table 13: Feature Importance for Escherichia coli - cefotaxime

| Feature Name | Gini Importance | Mutation Annotation | Gene |
|---|---|---|---|
| `0096HECL_ONPLLBGJ_03908_Endoribonuclease_HigB` | 0.009 | | |
| `'4483310','A:G','snp'` | 0.009 | synonymous_variant c.528T>C p.Asp176Asp | valS |
| `'2982569','G:A','snp'` | 0.008 | synonymous_variant c.690C>T p.Phe230Phe | kduD |
| `'4013889','C:T','snp'` | 0.008 | synonymous_variant c.837C>T p.Arg279Arg | metE |
| `'4231908','TTTT:CTTC','complex'` | 0.007 | synonymous_variant c.1323_1326delAAAAinsGAAG p.443 | lysC |
| `'2807452','G:A','snp'` | 0.007 | synonymous_variant c.321G>A p.Gly107Gly | proX |

**Table 13 – continued from previous page**

| Feature Name | Gini Importance | Mutation Annotation | Gene |
|---|---|---|---|
| '4430232','G:T','snp' | 0.005 | synonymous_variant c.369G>T p.Thr123Thr | cycA |
| '1512011','A:G','snp' | 0.005 | missense_variant c.358A>G p.Thr120Ala | ydcS |
| '576547','G:A','snp' | 0.005 | synonymous_variant c.279C>T p.Ser93Ser | nmpC |
| OHTPQWCQ_AKNJJMFD_04481_putative_RNA_guided_DNA_endonuclease_InsQ | 0.005 | | |

#### Escherichia coli - cefuroxime

| Bacteria | Antibiotic | F1 Score | MCC |
|---|---|---|---|
| Escherichia coli | cefuroxime | 0.506 | 0.440 |

Table 14: Feature Importance for Escherichia coli - cefuroxime

| Feature Name | Gini Importance | Mutation Annotation | Gene |
|---|---|---|---|
| 026HVE8R_CHODMJKP_03716_Beta_lactamase_CTX_M_1 | 0.005 | | |
| '3986737','A:C','snp' | 0.004 | synonymous_variant c.1146T>G p.Ala382Ala | hemY |
| '3103759','T:C','snp' | 0.004 | synonymous_variant c.747T>C p.Arg249Arg | mutY |
| '3799307','G:A','snp' | 0.004 | synonymous_variant c.6C>T p.Arg2Arg | waaU |
| RARXTY96_OCIINFJC_04553_Beta_lactamase_OXA_1 | 0.004 | | |
| '2339162','CATAG:TATAA','complex' | 0.003 | missense_variant c.255_259delCTATGinsTTATA p.Asp87Asn | gyrA |
| '2228922','A:G','snp' | 0.003 | | |
| '4125485','A:T','snp' | 0.003 | synonymous_variant c.1326T>A p.Ile442Ile | priA |
| '2226466','GACCA:TACCC','complex' | 0.003 | | |
| '1584606','G:A','snp' | 0.003 | synonymous_variant c.1881C>T p.Ser627Ser | ydeP |

#### Escherichia coli - trimethoprim_sulfamethoxazole

| Bacteria | Antibiotic | F1 Score | MCC |
|---|---|---|---|
| Escherichia coli | trimethoprim_sulfamethoxazole | 0.825 | 0.465 |

Table 15: Feature Importance for Escherichia coli - trimethoprim_sulfamethoxazole

| Feature Name | Gini Importance | Mutation Annotation | Gene |
|---|---|---|---|
| 0096HECL_ONPLLBGJ_04539_dihydropteroate_synthase | 0.011 | | |
| BUGAON43_PPMDDFLC_04867_hypothetical_protein | 0.008 | | |
| 3728VWZS_OHNGMDJJ_04695_hypothetical_protein | 0.008 | | |
| URIFEMFY_HMLGKCFH_04814_IS6_family_transposase_IS6100 | 0.004 | | |
| 0096HECL_ONPLLBGJ_04541_putative_chromate_transport_protein | 0.003 | | |
| 0096HECL_ONPLLBGJ_04545_Multidrug_efflux_pump_Tap | 0.003 | | |
| 5HCW7JCK_IOKNGGDD_04413_hypothetical_protein | 0.003 | | |
| ZYAAFH39_PPMCLIDB_03621_hypothetical_protein | 0.003 | | |
| IKMM4UV0_KIPEHDCH_04608_hypothetical_protein | 0.003 | | |
| 0096HECL_ONPLLBGJ_04611_Aminoglycoside_3__phosphotransferase | 0.003 | | |

#### Escherichia coli - ampicillin

| Bacteria | Antibiotic | F1 Score | MCC |
|---|---|---|---|
| Escherichia coli | ampicillin | 0.907 | 0.571 |

Table 16: Feature Importance for Escherichia coli - ampicillin

| Feature Name | Gini Importance | Mutation Annotation | Gene |
|---|---|---|---|
| 0096HECL_ONPLLBGJ_04660_Beta_lactamase_TEM | 0.011 | | |
| '3134461','C:T','snp' | 0.005 | synonymous_variant c.331C>T p.Leu111Leu | |

**Table 16 – continued from previous page**

| Feature Name | Gini Importance | Mutation Annotation | Gene |
|---|---|---|---|
| 016JET8K_BMHFLKKD_05009_hypothetical_protein | 0.004 | | |
| '4124948','C:T','snp' | 0.004 | synonymous_variant c.1863G>A p.Ala621Ala | priA |
| IR3P0GVX_HGMMIGBK_04679_multidrug_betaine_choline_efflux_transporter_EmrE | 0.004 | | |
| '2339173','G:A','snp' | 0.004 | missense_variant c.248C>T p.Ser83Leu | gyrA |
| '1958357','C:A','snp' | 0.003 | | |
| '2973437','A:C','snp' | 0.003 | synonymous_variant c.426T>G p.Ala142Ala | lplT |
| AWKE3LSI_PLPDHGMK_04566_Tn3_family_transposase_Tn2 | 0.003 | | |
| '3140407','T:G','snp' | 0.003 | synonymous_variant c.393A>C p.Ala131Ala | hybE |

#### Neisseria gonorrhoeae - azithromycin

| Bacteria | Antibiotic | F1 Score | MCC |
|---|---|---|---|
| Neisseria gonorrhoeae | azithromycin | 0.854 | 0.704 |

Table 17: Feature Importance for Neisseria gonorrhoeae - azithromycin

| Feature Name | Gini Importance | Mutation Annotation | Gene |
|---|---|---|---|
| '1739261','CCCCC:AAA','complex' | 0.013 | frameshift_variant&missense_variant c.681_685delCCCCCinsAAA p.Pro228fs | |
| '2065819','CGGAG:TGGAA','complex' | 0.007 | missense_variant c.153_157delCTCCGinsTTCCA p.Gly53Ser | |
| '166980','T:C','snp' | 0.007 | synonymous_variant c.1644T>C p.Leu548Leu | pepN |
| '2033171','CT:GC','complex' | 0.006 | missense_variant c.644_645delAGinsGC p.Glu215Gly | |
| '55341','CTAC:TTAG','complex' | 0.006 | missense_variant c.517_520delGTAGinsCTAA p.ValGlu173LeuLys | pilC |
| '1296502','C:A','snp' | 0.006 | missense_variant c.444G>T p.Glu148Asp | |
| '77410','G:A','snp' | 0.005 | synonymous_variant c.306G>A p.Val102Val | ispH |

Table 17 – continued from previous page

| Feature Name | Gini Importance | Mutation Annotation | | Gene |
|---|---|---|---|---|
| '2009329',' GCGGAAATGTG:AAAA',' complex' | 0.005 | | | |
| '816997','A:C','snp' | 0.005 | synonymous_variant p.Thr256Thr | c.768T>G | |
| '244219','A:C','snp' | 0.004 | | | |

#### Pseudomonas aeruginosa - levofloxacin

| Bacteria | Antibiotic | F1 Score | MCC |
|---|---|---|---|
| Pseudomonas aeruginosa | levofloxacin | 0.712 | 0.485 |

Table 18: Feature Importance for Pseudomonas aeruginosa - levofloxacin

| Feature Name | Gini Importance | Mutation Annotation | | Gene |
|---|---|---|---|---|
| '3558951','G:A','snp' | 0.009 | missense_variant p.Thr83Ile | c.248C>T | gyrA |
| 0OMPV1ZK_JIDMGLPJ_05713_site_specific_tyrosine_recombinase_XerD | 0.004 | | | |
| '465674','G:A','snp' | 0.003 | missense_variant p.Leu104Phe | c.310C>T | |
| '193820','G:T','snp' | 0.002 | synonymous_variant p.Ala120Ala | c.360C>A | |
| 0A28E4EJ_OOGCKHGP_02927_hypothetical_protein | 0.002 | | | |
| 4BDJYXUJ_KCCGMNJN_03014_dihydropteroate_synthase | 0.002 | | | |
| '4084660','G:A','snp' | 0.002 | synonymous_variant p.Leu118Leu | c.352C>T | |
| '4150304','G:A','snp' | 0.002 | synonymous_variant p.Val136Val | c.408C>T | |
| '1767798','C:T','snp' | 0.002 | | | |
| '1942753','A:G','snp' | 0.002 | synonymous_variant p.Thr64Thr | c.192T>C | ppiB |

#### Pseudomonas aeruginosa - imipenem

| Bacteria | Antibiotic | F1 Score | MCC |
|---|---|---|---|
| Pseudomonas aeruginosa | imipenem | 0.607 | 0.331 |

Table 19: Feature Importance for Pseudomonas aeruginosa - imipenem

| Feature Name | Gini Importance | Mutation Annotation | Gene |
|---|---|---|---|
| 0OMPV1ZK_JIDMGLPJ_05713_site_specific_tyrosine_recombinase_XerD | 0.003 | | |
| 0A28E4EJ_OOGCKHGP_02968_hypothetical_protein | 0.003 | | |
| 0EQSDK2Y_NCFANGKH_03221_Tyrosine_recombinase_XerC | 0.002 | | |
| '6188056','G:T','snp' | 0.002 | | |
| '5140761','GCCT:CCCA','complex' | 0.002 | synonymous_variant c.426_429delAGGCinsTGGG p.144 | pra |
| '3558951','G:A','snp' | 0.002 | missense_variant c.248C>T p.Thr83Ile | gyrA |
| '2214301','C:T','snp' | 0.002 | synonymous_variant c.609C>T p.Arg203Arg | |
| '4392709','T:C','snp' | 0.002 | synonymous_variant c.1653A>G p.Ala551Ala | |
| '4825342','G:A','snp' | 0.002 | synonymous_variant c.585C>T p.Gly195Gly | tadB |
| 0349MMD9_MKKACBFB_05411_hypothetical_protein | 0.002 | | |

#### Pseudomonas aeruginosa - amikacin

| Bacteria | Antibiotic | F1 Score | MCC |
|---|---|---|---|
| Pseudomonas aeruginosa | amikacin | 0.500 | 0.403 |

Table 20: Feature Importance for Pseudomonas aeruginosa - amikacin

| Feature Name | Gini Importance | Mutation Annotation | Gene |
|---|---|---|---|
| 4BDJYXUJ_KCCGMNJN_03014_dihydropteroate_synthase | 0.004 | | |
| 0NWCMZDU_HHCALAAC_03374_hypothetical_protein | 0.002 | | |
| '3256275','T:C','snp' | 0.002 | missense_variant c.337A>G p.Met113Val | |
| 2QQ1LXQV_EMIGCGFH_06414_hypothetical_protein | 0.002 | | |

**Table 20 – continued from previous page**

| Feature Name | Gini Importance | Mutation Annotation | Gene |
|---|---|---|---|
| '1771737','TGGTCGGGC:GGGCCGGAT','complex' | 0.002 | missense_variant c.907_915delGCCCGACCAinsATCCGGCCC p.Ala303Ile | |
| 1LDUFUPH_DONMHOKI_06032_hypothetical_protein | 0.002 | | |
| '1791286','C:A','snp' | 0.002 | missense_variant c.207C>A p.Asp69Glu | |
| 0EQSDK2Y_NCFANGKH_01931_beta_lactamase | 0.002 | | |
| '3475560','G:A','snp' | 0.002 | | |
| 0NWCMZDU_HHCALAAC_03370_hypothetical_protein | 0.002 | | |

#### Pseudomonas aeruginosa - ciprofloxacin

| Bacteria | Antibiotic | F1 Score | MCC |
|---|---|---|---|
| Pseudomonas aeruginosa | ciprofloxacin | 0.591 | 0.403 |

Table 21: Feature Importance for Pseudomonas aeruginosa - ciprofloxacin

| Feature Name | Gini Importance | Mutation Annotation | Gene |
|---|---|---|---|
| '3558951','G:A','snp' | 0.010 | missense_variant c.248C>T p.Thr83Ile | gyrA |
| '4472763','T:C','snp' | 0.002 | synonymous_variant c.633T>C p.Ile211Ile | |
| 0OMPV1ZK_JIDMGLPJ_00734_glycine_betaine_transmethylase | 0.002 | | |
| '4465457','A:T','snp' | 0.002 | | |
| '6231643','A:G','snp' | 0.002 | | |
| '5735248','G:C','snp' | 0.002 | synonymous_variant c.399C>G p.Gly133Gly | |
| '3488264','A:G','snp' | 0.002 | synonymous_variant c.129T>C p.Thr43Thr | metZ |
| 0OMPV1ZK_JIDMGLPJ_01245_hypothetical_protein | 0.002 | | |
| '4140771','T:C','snp' | 0.002 | | |
| '1672235','G:A','snp' | 0.002 | synonymous_variant c.156G>A p.Lys52Lys | |

#### Pseudomonas aeruginosa - ceftazidime

| Bacteria | Antibiotic | F1 Score | MCC |
|---|---|---|---|
| Pseudomonas aeruginosa | ceftazidime | 0.389 | 0.196 |

Table 22: Feature Importance for Pseudomonas aeruginosa - ceftazidime

| Feature Name | Gini Importance | Mutation Annotation | Gene |
|---|---|---|---|
| '570858','T:TG','ins' | 0.003 | | |
| '3275668','G:C','snp' | 0.003 | synonymous_variant c.567C>G p.Arg189Arg | |
| OOMPV1ZK_JIDMGLPJ_05338_hypothetical_protein | 0.002 | | |
| '5156629','G:A','snp' | 0.002 | missense_variant c.262G>A p.Val88Met | morA |
| '3442046','T:G','snp' | 0.002 | synonymous_variant c.438A>C p.Ile146Ile | |
| '3957297','C:T','snp' | 0.002 | synonymous_variant c.2391C>T p.Gly797Gly | |
| '4118102','G:A','snp' | 0.002 | synonymous_variant c.1164C>T p.Ile388Ile | |
| '1404724','G:A','snp' | 0.002 | synonymous_variant c.474C>T p.Tyr158Tyr | |
| '5293981','A:G','snp' | 0.002 | synonymous_variant c.1020T>C p.His340His | |
| '4984310','T:C','snp' | 0.002 | synonymous_variant c.1161A>G p.Glu387Glu | murA |

#### Pseudomonas aeruginosa - cefepime

| Bacteria | Antibiotic | F1 Score | MCC |
|---|---|---|---|
| Pseudomonas aeruginosa | cefepime | 0.667 | 0.350 |

Table 23: Feature Importance for Pseudomonas aeruginosa - cefepime

| Feature Name | Gini Importance | Mutation Annotation | Gene |
|---|---|---|---|
| '3584487','A:G','snp' | 0.003 | synonymous_variant c.888T>C p.Thr296Thr | glk |
| '4188555','C:T','snp' | 0.003 | synonymous_variant c.993G>A p.Ala331Ala | hom |
| '4326625','A:AC','ins' | 0.002 | | |
| '6072431','A:G','snp' | 0.002 | synonymous_variant c.954T>C p.Gly318Gly | cls |
| '4722547','A:G','snp' | 0.002 | synonymous_variant c.312T>C p.Ser104Ser | |

**Table 23 – continued from previous page**

| Feature Name | Gini Importance | Mutation Annotation | Gene |
|---|---|---|---|
| '437196','A:G','snp' | 0.002 | synonymous_variant c.627A>G p.Glu209Glu | pilT |
| '4177057','T:C','snp' | 0.002 | synonymous_variant c.2070A>G p.Gly690Gly | |
| '2868219','A:T','snp' | 0.002 | missense_variant c.1084T>A p.Cys362Ser | |
| '3172933','T:C','snp' | 0.002 | synonymous_variant c.243A>G p.Leu81Leu | arr |
| '2946204','G:A','snp' | 0.002 | synonymous_variant c.339G>A p.Pro113Pro | |

#### Pseudomonas aeruginosa - meropenem

| Bacteria | Antibiotic | F1 Score | MCC |
|---|---|---|---|
| Pseudomonas aeruginosa | meropenem | 0.619 | 0.246 |

Table 24: Feature Importance for Pseudomonas aeruginosa - meropenem

| Feature Name | Gini Importance | Mutation Annotation | Gene |
|---|---|---|---|
| 4BDJYXUJ_KCCGMNJN_03014_dihydropteroate_synthase | 0.003 | | |
| 0OMPV1ZK_JIDMGLPJ_03649_hypothetical_protein | 0.002 | | |
| 0OMPV1ZK_JIDMGLPJ_03623_hypothetical_protein | 0.002 | | |
| 6BVE2G1L_PMJBMGCC_01182_porin_D | 0.002 | | |
| '5574227','G:A','snp' | 0.002 | missense_variant c.260C>T p.Ser87Leu | parC |
| 02UYU1WT_KIFDEIGC_03338_hypothetical_protein | 0.002 | | |
| 0349MMD9_MKKACBFB_03573_IS6_family_transposase_IS6100 | 0.002 | | |
| 0349MMD9_MKKACBFB_05411_hypothetical_protein | 0.002 | | |
| 0OMPV1ZK_JIDMGLPJ_05717_hypothetical_protein | 0.002 | | |
| '2100789','T:C','snp' | 0.001 | synonymous_variant c.1338T>C p.Asp446Asp | |

#### Staphylococcus aureus- tetracycline

| Bacteria | Antibiotic | F1 Score | MCC |
|---|---|---|---|
| Staphylococcus aureus | tetracycline | 0.879 | 0.859 |

Table 25: Feature Importance for Staphylococcus aureus- tetracycline

| Feature Name | Gini Importance | Mutation Annotation | Gene |
|---|---|---|---|
| '1679902','G:A','snp' | 0.011 | synonymous_variant c.687C>T p.Ala229Ala | tig |
| 05L2U7U6_NAKNMPGO_02803_dihydrolipoamide_dehydrogenase | 0.011 | | |
| '854310','G:T','snp' | 0.010 | missense_variant c.162C>A p.Asp54Glu | |
| '428810','T:C','snp' | 0.010 | synonymous_variant c.258A>G p.Lys86Lys | |
| '791830','C:A','snp' | 0.010 | missense_variant c.298C>A p.His100Asn | |
| 05L2U7U6_NAKNMPGO_02839_tetracycline_efflux_MFS_transporter_Tet_K | 0.007 | | |
| '699570','T:C','snp' | 0.006 | missense_variant c.680A>G p.Asn227Ser | |
| 05L2U7U6_NAKNMPGO_02463_hypothetical_protein | 0.006 | | |
| '1189093','T:C','snp' | 0.006 | synonymous_variant c.1536T>C p.Val512Val | |
| '615798','T:C','snp' | 0.006 | synonymous_variant c.870T>C p.Ile290Ile | |

#### Staphylococcus aureus - trimethoprim_sulfamethoxazole

| Bacteria | Antibiotic | F1 Score | MCC |
|---|---|---|---|
| Staphylococcus aureus | trimethoprim_sulfamethoxazole | 0.875 | 0.840 |

Table 26: Feature Importance for Staphylococcus aureus- trimethoprim_sulfamethoxazole

| Feature Name | Gini Importance | Mutation Annotation | Gene |
|---|---|---|---|
| '308557','G:A','snp' | 0.019 | synonymous_variant c.87C>T p.Thr29Thr | |
| '14584','TGAC:CGAT','complex' | 0.014 | | |

**Table 26 – continued from previous page**

| Feature Name | Gini Importance | Mutation Annotation | Gene |
|---|---|---|---|
| '14621','TGAT: CGGA','complex' | 0.014 | | |
| '2687872','C:A','snp' | 0.014 | | |
| '158446','G:C','snp' | 0.014 | missense_variant c.273C>G p.His91Gln | |
| '148841','G:T','snp' | 0.014 | | |
| '77496','T:C','snp' | 0.013 | | |
| '2744748','CACT: TACG','complex' | 0.013 | synonymous_variant c.480_483delCACTinsTACG p.162 | |
| '77265','TAAT: CAAG','complex' | 0.013 | synonymous_variant c.168_171delATTAinsCTTG p.58 | |
| '100183','A:G','snp' | 0.013 | | |

#### Staphylococcus aureus - clindamycin

| Bacteria | Antibiotic | F1 Score | MCC |
|---|---|---|---|
| Staphylococcus aureus | clindamycin | 0.927 | 0.884 |

Table 27: Feature Importance for Staphylococcus aureus - clindamycin

| Feature Name | Gini Importance | Mutation Annotation | Gene |
|---|---|---|---|
| '324225','G:A','snp' | 0.016 | synonymous_variant c.768G>A p.Leu256Leu | |
| '250761','CGAC: TGAA','complex' | 0.011 | missense_variant c.171_174delCGACinsTGAA p.Asp58Glu | |
| OOFM2TYT_ NNALBEHK_02547_ Glycerophosphodiester_ phosphodiesterase_ _cytoplasmic | 0.011 | | |
| '139673','T:C','snp' | 0.010 | synonymous_variant c.273T>C p.Phe91Phe | |
| 0130GIA4_ MKGLEHDE_02493_ dimethyladenosine_ transferase | 0.009 | | |
| '2734812','ATGAA: GTGAT','complex' | 0.009 | synonymous_variant c.339_343delTTCATinsATCAC p.116 | |
| '2686892','T:C','snp' | 0.009 | synonymous_variant c.414A>G p.Leu138Leu | |
| '98260','A:C','snp' | 0.008 | synonymous_variant c.471A>C p.Val157Val | |
| '2771331','G:A','snp' | 0.008 | | |

**Table 27 – continued from previous page**

| Feature Name | Gini Importance | Mutation Annotation | Gene |
|---|---|---|---|
| '7255','C:T','snp' | 0.008 | missense_variant c.251C>T p.Ser84Leu | |

#### Staphylococcus aureus - ciprofloxacin

| Bacteria | Antibiotic | F1 Score | MCC |
|---|---|---|---|
| Staphylococcus aureus | ciprofloxacin | 0.931 | 0.886 |

Table 28: Feature Importance for Staphylococcus aureus - ciprofloxacin

| Feature Name | Gini Importance | Mutation Annotation | Gene |
|---|---|---|---|
| '2640119','A:G','snp' | 0.010 | synonymous_variant c.990T>C p.Asn330Asn | |
| '2537427','G:A','snp' | 0.009 | synonymous_variant c.117G>A p.Pro39Pro | |
| 185UP9BH_FCBKHAEB_02529_hypothetical_protein | 0.008 | | |
| '649607','T:C','snp' | 0.008 | | |
| 00FM2TYT_NNALBEHK_02549_PBP2a_family_beta_lactam_resistant_peptidoglycan_transpeptidase_MecA | 0.007 | | |
| '1440199','A:G','snp' | 0.007 | synonymous_variant c.186T>C p.Val62Val | aroB |
| 00FM2TYT_NNALBEHK_02548_hypothetical_protein | 0.006 | | |
| 120AWVQ8_FPALCBMI_00835_hypothetical_protein | 0.006 | | |
| '107745','TGGA:CGGT','complex' | 0.006 | synonymous_variant c.372_375delTCCAinsACCG p.126 | |
| '2502773','C:T','snp' | 0.006 | | |

#### Staphylococcus aureus - methicillin

| Bacteria | Antibiotic | F1 Score | MCC |
|---|---|---|---|
| Staphylococcus aureus | methicillin | 0.933 | 0.879 |

Table 29: Feature Importance for Staphylococcus aureus - methicillin

| Feature Name | Gini Importance | Mutation Annotation | Gene |
|---|---|---|---|
| 00FM2TYT_NNALBEHK_02548_hypothetical_protein | 0.015 | | |
| 00FM2TYT_NNALBEHK_02226_hypothetical_protein | 0.013 | | |
| '2010091','A:G','snp' | 0.011 | synonymous_variant c.1200T>C p.Val400Val | |
| '2495653','T:C','snp' | 0.011 | synonymous_variant c.333A>G p.Gly111Gly | |
| '199266','G:A','snp' | 0.010 | | |
| '2025570','C:T','snp' | 0.009 | synonymous_variant c.765G>A p.Ala255Ala | |
| 00FM2TYT_NNALBEHK_02549_PBP2a_family_beta_lactam_resistant_peptidoglycan_transpeptidase_MecA | 0.007 | | |
| '677055','C:A','snp' | 0.007 | missense_variant c.1153C>A p.Gln385Lys | |
| '395419','T:C','snp' | 0.006 | synonymous_variant c.183T>C p.Ser61Ser | |
| '1120371','G:A','snp' | 0.005 | synonymous_variant c.1422G>A p.Glu474Glu | carB |

#### Staphylococcus aureus - oxacillin

| Bacteria | Antibiotic | F1 Score | MCC |
|---|---|---|---|
| Staphylococcus aureus | oxacillin | 0.907 | 0.863 |

Table 30: Feature Importance for Staphylococcus aureus - oxacillin

| Feature Name | Gini Importance | Mutation Annotation | Gene |
|---|---|---|---|
| '1525642','T:C','snp' | 0.015 | synonymous_variant c.408T>C p.Pro136Pro | |
| '440860','C:T','snp' | 0.014 | synonymous_variant c.717C>T p.Tyr239Tyr | |
| '2313997','A:G','snp' | 0.014 | synonymous_variant c.603T>C p.Ile201Ile | rpsC |
| '2476208','A:G','snp' | 0.014 | missense_variant c.679T>C p.Cys227Arg | |
| '21536','A:T','snp' | 0.012 | synonymous_variant c.771A>T p.Gly257Gly | |
| '1828726','C:T','snp' | 0.011 | synonymous_variant c.258C>T p.Ser86Ser | |

**Table 30 – continued from previous page**

| Feature Name | Gini Importance | Mutation Annotation | Gene |
|---|---|---|---|
| '411316','A:G','snp' | 0.008 | missense_variant c.595A>G p.Met199Val | |
| '1353284','A:G','snp' | 0.008 | synonymous_variant c.27T>C p.Asn9Asn | |
| '702109','A:G','snp' | 0.008 | synonymous_variant c.57T>C p.Gly19Gly | |
| '1130174','A:G','snp' | 0.008 | synonymous_variant c.1068A>G p.Glu356Glu | |

#### Staphylococcus aureus - gentamicin

| Bacteria | Antibiotic | F1 Score | MCC |
|---|---|---|---|
| Staphylococcus aureus | gentamicin | 0.930 | 0.917 |

Table 31: Feature Importance for Staphylococcus aureus - gentamicin

| Feature Name | Gini Importance | Mutation Annotation | Gene |
|---|---|---|---|
| '1525570','C:T','snp' | 0.023 | synonymous_variant c.336C>T p.Ala112Ala | |
| '2419368','A:G','snp' | 0.015 | synonymous_variant c.75T>C p.Phe25Phe | |
| BR8Z1NIG_IIKKHCGP_02633_Bifunctional_AAC_APH | 0.015 | | |
| '744269','A:G','snp' | 0.015 | | |
| '2595499','T:C','snp' | 0.015 | synonymous_variant c.174T>C p.Thr58Thr | |
| '602943','A:T','snp' | 0.014 | synonymous_variant c.978A>T p.Val326Val | argS |
| '1703357','C:T','snp' | 0.014 | synonymous_variant c.312G>A p.Thr104Thr | |
| '1579166','C:T','snp' | 0.014 | | |
| '973544','T:A','snp' | 0.014 | synonymous_variant c.1923A>T p.Ala641Ala | |
| '2431382','A:G','snp' | 0.014 | synonymous_variant c.1248T>C p.Val416Val | |

#### Staphylococcus aureus - erythromycin

| Bacteria | Antibiotic | F1 Score | MCC |
|---|---|---|---|
| Staphylococcus aureus | erythromycin | 0.821 | 0.718 |

Table 32: Feature Importance for Staphylococcus aureus - erythromycin

| Feature Name | Gini Importance | Mutation Annotation | Gene |
|---|---|---|---|
| 00FM2TYT_NNALBEHK_02547_Glycerophosphodiester_phosphodiesterase__cytoplasmic | 0.016 | | |
| 0550Z5VB_NEAFNFFJ_02708_beta_lactam_sensor_signal_transducer_MecR1 | 0.009 | | |
| 0130GIA4_MKGLEHDE_02493_dimethyladenosine_transferase | 0.008 | | |
| '270896','T:C','snp' | 0.008 | missense_variant c.616A>G p.Thr206Ala | |
| 0130GIA4_MKGLEHDE_02494_hypothetical_protein | 0.008 | | |
| '1911743','C:T','snp' | 0.007 | synonymous_variant c.852C>T p.Ser284Ser | |
| '2599453','A:G','snp' | 0.006 | synonymous_variant c.675A>G p.Thr225Thr | |
| '2690096','T:G','snp' | 0.006 | synonymous_variant c.763A>C p.Arg255Arg | |
| '1193878','C:T','snp' | 0.006 | | |
| '1380915','A:G','snp' | 0.006 | synonymous_variant c.23784T>C p.Asn7928Asn | |

#### Staphylococcus aureus - penicillin

| Bacteria | Antibiotic | F1 Score | MCC |
|---|---|---|---|
| Staphylococcus aureus | penicillin | 0.948 | 0.606 |

Table 33: Feature Importance for Staphylococcus aureus - penicillin

| Feature Name | Gini Importance | Mutation Annotation | Gene |
|---|---|---|---|
| 0EXX2GFC_OCNDEOAA_02626_beta_lactam_sensor_signal_transducer_BlaR1 | 0.010 | | |

**Table 33 – continued from previous page**

| Feature Name | Gini Importance | Mutation Annotation | Gene |
|---|---|---|---|
| 01LDN2PD_FIAEPMAC_02591_Cadmium_resistance_transcriptional_regulatory_protein_CadC | 0.008 | | |
| 0550Z5VB_NEAFNFFJ_02675_Plasmid_recombination_enzyme | 0.008 | | |
| 002GHROQ_IILPEGAA_01060_beta_lactam_sensor_signal_transducer_BlaR1 | 0.008 | | |
| 01LDN2PD_FIAEPMAC_02574_DNA_invertase_hin | 0.007 | | |
| 0D4NEYJ9_MABFDMMK_02667_hypothetical_protein | 0.007 | | |
| 71AL04U1_LECODONG_02582_hypothetical_protein | 0.006 | | |
| 0130GIA4_MKGLEHDE_02348_Beta_lactamase | 0.006 | | |
| 01LDN2PD_FIAEPMAC_02604_hypothetical_protein | 0.005 | | |
| '1489551','T:C','snp' | 0.005 | synonymous_variant c.858A>G p.Gln286Gln | |

#### Streptococcus pneumoniae - chloramphenicol

| Bacteria | Antibiotic | F1 Score | MCC |
|---|---|---|---|
| Streptococcus pneumoniae | chloramphenicol | 0.933 | 0.911 |

Table 34: Feature Importance for Streptococcus pneumoniae - chloramphenicol

| Feature Name | Gini Importance | Mutation Annotation | Gene |
|---|---|---|---|
| 005GYZ04_IGMBNOGL_00106_conjugal_transfer_protein_TrbL | 0.018 | | |
| '1102372','T:C','snp' | 0.016 | synonymous_variant c.3936T>C p.Gly1312Gly | |
| '1100809','T:C','snp' | 0.016 | synonymous_variant c.2373T>C p.Gly791Gly | |
| '1099716','A:G','snp' | 0.016 | missense_variant c.1280A>G p.Gln427Arg | |

**Table 34 – continued from previous page**

| Feature Name | Gini Importance | Mutation Annotation | Gene |
|---|---|---|---|
| '1093394','A:G','snp' | 0.016 | synonymous_variant c.1563A>G p.Glu521Glu | |
| '883755','C:A','snp' | 0.015 | synonymous_variant c.357G>T p.Thr119Thr | |
| '1091411','G:A','snp' | 0.015 | synonymous_variant c.798G>A p.Leu266Leu | |
| '1941166','G:A','snp' | 0.015 | synonymous_variant c.1710C>T p.Val570Val | |
| '1104730','TA:CG','complex' | 0.014 | | |
| '1111372','C:T','snp' | 0.014 | synonymous_variant c.180C>T p.Pro60Pro | |

#### Streptococcus pneumoniae - erythromycin

| Bacteria | Antibiotic | F1 Score | MCC |
|---|---|---|---|
| Streptococcus pneumoniae | erythromycin | 0.882 | 0.751 |

Table 35: Feature Importance for Streptococcus pneumoniae - erythromycin

| Feature Name | Gini Importance | Mutation Annotation | Gene |
|---|---|---|---|
| 02OM9Y49_ADCIONGJ_00150_hypothetical_protein | 0.013 | | |
| 02OM9Y49_ADCIONGJ_00153_ABC_F_type_ribosomal_protection_protein_Msr_D | 0.012 | | |
| 02OM9Y49_ADCIONGJ_00155_hypothetical_protein | 0.012 | | |
| '1199618','T:C','snp' | 0.007 | missense_variant c.437A>G p.Asn146Ser | |
| '1246257','AC:GT','complex' | 0.006 | missense_variant c.668_669delGTinsAC p.Ser223Asn | |
| 005GYZ04_IGMBNOGL_00136_ATP_binding_protein | 0.005 | | |
| '1789997','A:C','snp' | 0.005 | missense_variant c.1121T>G p.Val374Gly | |
| '1532651','AC:GT','complex' | 0.005 | synonymous_variant c.33_34delACinsGT p.13 | |
| '1530636','CTCT:TTCA','complex' | 0.005 | synonymous_variant c.1848_1851delAGAGinsTGAA p.618 | |

**Table 35 – continued from previous page**

| Feature Name | Gini Importance | Mutation Annotation | Gene |
|---|---|---|---|
| '351417','A:G','snp' | 0.005 | missense_variant c.11A>G p.Lys4Arg | ftsL |

#### Streptococcus pneumoniae - penicillin

| Bacteria | Antibiotic | F1 Score | MCC |
|---|---|---|---|
| Streptococcus pneumoniae | penicillin | 0.913 | 0.886 |

Table 36: Feature Importance for Streptococcus pneumoniae - penicillin

| Feature Name | Gini Importance | Mutation Annotation | Gene |
|---|---|---|---|
| '1579849','C:T','snp' | 0.021 | | |
| '1567084','T:C','snp' | 0.017 | missense_variant c.706A>G p.Thr236Ala | divIVA |
| '1578191','A:T','snp' | 0.016 | missense_variant c.1480T>A p.Ser494Thr | pbp2b |
| '1868464','A:G','snp' | 0.013 | synonymous_variant c.102A>G p.Arg34Arg | |
| '540730','A:G','snp' | 0.011 | synonymous_variant c.1245T>C p.Ile415Ile | |
| '1702579','C:T','snp' | 0.011 | missense_variant c.454G>A p.Ala152Thr | |
| '1266451','GAAAGTT: AAAGGTG','complex' | 0.011 | synonymous_variant c.423_429delGAAAGTTinsAAAGGTG p.144 | |
| '509378','CACG: TACA','complex' | 0.011 | | |
| '1578207','A:G','snp' | 0.010 | synonymous_variant c.1464T>C p.Gly488Gly | pbp2b |
| '1578301','A:G','snp' | 0.009 | missense_variant c.1370T>C p.Val457Ala | pbp2b |

#### Streptococcus pneumoniae - trimethoprim_sulfamethoxazole

| Bacteria | Antibiotic | F1 Score | MCC |
|---|---|---|---|
| Streptococcus pneumoniae | trimethoprim_sulfamethoxazole | 0.952 | 0.835 |

**Table 37 – continued from previous page**

| Feature Name | Gini Importance | Mutation Annotation | Gene |
|---|---|---|---|

Table 37: Feature Importance for Streptococcus pneumoniae - trimethoprim_sulfamethoxazole

| Feature Name | Gini Importance | Mutation Annotation | Gene |
|---|---|---|---|
| '317350','GGCT: AGCC','complex' | 0.011 | synonymous_variant c.168_171delGGCTinsAGCC p.58 | folP |
| '1490669','C:T','snp' | 0.008 | synonymous_variant c.378G>A p.Glu126Glu | |
| '1490735','C:T','snp' | 0.008 | synonymous_variant c.312G>A p.Gln104Gln | |
| '1490505',' CATGGCTGTAAAAAA: AAAAGCCGTGAATAG',' complex' | 0.007 | missense_variant c.22_36delTTTTTTACAGCCATGinsCTATTCACGGCTTTT p.PhePheThrAlaMet8LeuPheThrAlaPhe | |
| '317362','A:C','snp' | 0.007 | synonymous_variant c.180A>C p.Leu60Leu | folP |
| '1490717','C:T','snp' | 0.006 | synonymous_variant c.330G>A p.Glu110Glu | |
| '1490749','GAATA: TAATG','complex' | 0.005 | missense_variant c.294_298delTATTCinsCATTA p.Leu100Ile | |
| 008F7RZS_CEKMKMCL_ 01618_dihydrofolate_ reductase | 0.005 | | |
| '1578081','TGGG: CGGC','complex' | 0.005 | synonymous_variant c.1587_1590delCCCAinsGCCG p.531 | pbp2b |
| '1490846','T:C','snp' | 0.005 | synonymous_variant c.201A>G p.Thr67Thr | |

#### Streptococcus pneumoniae - tetracycline

| Bacteria | Antibiotic | F1 Score | MCC |
|---|---|---|---|
| Streptococcus pneumoniae | tetracycline | 0.959 | 0.928 |

Table 38: Feature Importance for Streptococcus pneumoniae - tetracycline

| Feature Name | Gini Importance | Mutation Annotation | Gene |
|---|---|---|---|
| 0G05QLHE_PBBODDHP_ 00047_conjugal_ transfer_protein | 0.018 | | |

**Table 38 – continued from previous page**

| Feature Name | Gini Importance | Mutation Annotation | Gene |
|---|---|---|---|
| 94RH3ZSE_DJPOLNNL_00852_tyrosine_type_recombinase_integrase | 0.014 | | |
| '1251310','G:A','snp' | 0.014 | missense_variant c.100C>T p.Leu34Phe | |
| '2102277','T:C','snp' | 0.012 | missense_variant c.1253A>G p.Asn418Ser | hk06 |
| 005GYZO4_IGMBNOGL_00136_ATP_binding_protein | 0.010 | | |
| '1542357','G:T','snp' | 0.010 | missense_variant c.271C>A p.His91Asn | |
| 005GYZO4_IGMBNOGL_00135_hypothetical_protein | 0.010 | | |
| '316105','T:C','snp' | 0.009 | missense_variant c.1346T>C p.Val449Ala | |
| DN6ALO21_PHNEMFFI_00041_tetracycline_resistance_ribosomal_protection_protein_Tet_M | 0.009 | | |
| '58736','C:A','snp' | 0.009 | missense_variant c.1270C>A p.Gln424Lys | radA |

#### Peptoclostridium difficile - moxifloxacin

| Bacteria | Antibiotic | F1 Score | MCC |
|---|---|---|---|
| Peptoclostridium difficile | moxifloxacin | 0.931 | 0.883 |

Table 39: Feature Importance for Peptoclostridium difficile - moxifloxacin

| Feature Name | Gini Importance | Mutation Annotation | Gene |
|---|---|---|---|
| '2418315','A:AT','ins' | 0.021 | frameshift_variant c.532_533insA p.Phe178fs | |
| '3623297','TGGG:AGGA','complex' | 0.014 | synonymous_variant c.159_162delCCCAinsTCCT p.55 | |
| '3610229','A:G','snp' | 0.013 | synonymous_variant c.1065T>C p.Val355Val | |
| '778998','A:G','snp' | 0.013 | synonymous_variant c.168A>G p.Gly56Gly | |
| '1462303','G:A','snp' | 0.011 | missense_variant c.1015G>A p.Glu339Lys | |
| 11PRF2EA_NJHBEIBE_00383_hypothetical_protein | 0.009 | | |

**Table 39 – continued from previous page**

| Feature Name | Gini Importance | Mutation Annotation | Gene |
|---|---|---|---|
| '1135475','GA:G','del' | 0.008 | frameshift_variant c.753delA p.Ala252fs | ytvI |
| 0KHGBGW1_LPBNNBCB_02253_ABC_transporter_ATP_binding_protein | 0.008 | | |
| '3800458','A:G','snp' | 0.008 | synonymous_variant c.420T>C p.Ser140Ser | |
| '2455165','T:C','snp' | 0.008 | synonymous_variant c.135A>G p.Lys45Lys | |

#### Peptoclostridium difficile - azithromycin

| Bacteria | Antibiotic | F1 Score | MCC |
|---|---|---|---|
| Peptoclostridium difficile | azithromycin | 0.889 | 0.799 |

Table 40: Feature Importance for Peptoclostridium difficile - azithromycin

| Feature Name | Gini Importance | Mutation Annotation | Gene |
|---|---|---|---|
| '713045','C:T','snp' | 0.018 | synonymous_variant c.222C>T p.Ser74Ser | |
| 0KHGBGW1_LPBNNBCB_01782_hypothetical_protein | 0.013 | | |
| '2579272','G:A','snp' | 0.013 | synonymous_variant c.1086C>T p.Gly362Gly | |
| '2455069','ACCA:GTTT','complex' | 0.013 | missense_variant c.228_231delTGGTinsAAAC p.AspGly76GluAsn | |
| '3625075','T:C','snp' | 0.012 | missense_variant c.1396A>G p.Ile466Val | |
| 0KHGBGW1_LPBNNBCB_02254_hypothetical_protein | 0.012 | | |
| 0KHGBGW1_LPBNNBCB_00462_hypothetical_protein | 0.012 | | |
| '570890','G:A','snp' | 0.012 | missense_variant c.1108G>A p.Val370Ile | |
| 0MO5GOHY_KPLIKJPD_02345_GNAT_family_N_acetyltransferase | 0.008 | | |
| '703066','A:G','snp' | 0.007 | missense_variant c.998A>G p.Lys333Arg | |

#### Peptoclostridium difficile - clarithromycin

| Bacteria | Antibiotic | F1 Score | MCC |
|---|---|---|---|
| Peptoclostridium difficile | clarithromycin | 0.937 | 0.885 |

Table 41: Feature Importance for Peptoclostridium difficile - clarithromycin

| Feature Name | Gini Importance | Mutation Annotation | Gene |
|---|---|---|---|
| '2455165','T:C','snp' | 0.020 | synonymous_variant c.135A>G p.Lys45Lys | |
| '1451666','C:A','snp' | 0.013 | missense_variant c.719C>A p.Ala240Asp | |
| 0KHGBGW1_LPBNNBCB_02262_DNA_replication_and_repair_protein_RecF | 0.012 | | |
| '2531805','T:G','snp' | 0.012 | missense_variant c.205A>C p.Ile69Leu | nadA |
| '3625135','C:T','snp' | 0.012 | missense_variant c.1336G>A p.Ala446Thr | |
| '2330424','C:T','snp' | 0.010 | synonymous_variant c.237G>A p.Val79Val | |
| 0KHGBGW1_LPBNNBCB_02273_hypothetical_protein | 0.008 | | |
| 11PRF2EA_NJHBEIBE_00383_hypothetical_protein | 0.007 | | |
| 0KHGBGW1_LPBNNBCB_02271_hypothetical_protein | 0.007 | | |
| '931051','C:T','snp' | 0.007 | missense_variant c.31C>T p.Pro11Ser | |

#### Acinetobacter baumannii - ciprofloxacin

| Bacteria | Antibiotic | F1 Score | MCC |
|---|---|---|---|
| Acinetobacter baumannii | ciprofloxacin | 0.972 | 0.765 |

Table 42: Feature Importance for Acinetobacter baumannii - ciprofloxacin

| Feature Name | Gini Importance | Mutation Annotation | Gene |
|---|---|---|---|
| '2092345','C:T','snp' | 0.012 | synonymous_variant c.48C>T p.Ser16Ser | |
| '556270','C:T','snp' | 0.011 | synonymous_variant c.402C>T p.Gly134Gly | |

**Table 42 – continued from previous page**

| Feature Name | Gini Importance | Mutation Annotation | Gene |
|---|---|---|---|
| '2846948','CGCA:TGCG','complex' | 0.009 | synonymous_variant c.1956_1959delTGCGinsCGCA p.654 | |
| '190219','TGCT:GGCA','complex' | 0.006 | synonymous_variant c.1899_1902delTGCTinsGGCA p.635 | acs |
| '1420307','A:G','snp' | 0.006 | synonymous_variant c.288A>G p.Leu96Leu | |
| '1632631','A:G','snp' | 0.006 | synonymous_variant c.1239A>G p.Ala413Ala | |
| '1143964','C:T','snp' | 0.006 | | |
| '1691931','G:T','snp' | 0.005 | missense_variant c.34C>A p.Pro12Thr | |
| '2847212','ACTT:TTTC','complex' | 0.005 | missense_variant c.1692_1695delAAGTinsGAAA p.Ser565Lys | |
| '1267301','AAAAGGT:GAGAGGG','complex' | 0.005 | | |

#### Acinetobacter baumannii - gentamicin

| Bacteria | Antibiotic | F1 Score | MCC |
|---|---|---|---|
| Acinetobacter baumannii | gentamicin | 0.976 | 0.787 |

Table 43: Feature Importance for Acinetobacter baumannii - gentamicin

| Feature Name | Gini Importance | Mutation Annotation | Gene |
|---|---|---|---|
| '3131204','C:T','snp' | 0.010 | missense_variant c.737C>T p.Pro246Leu | |
| '1156932','A:G','snp' | 0.008 | synonymous_variant c.1143T>C p.Val381Val | |
| '187060','T:A','snp' | 0.008 | synonymous_variant c.279A>T p.Leu93Leu | |
| '3162915','A:T','snp' | 0.007 | synonymous_variant c.90T>A p.Ala30Ala | |
| '2969477','AGATGCT:GGAAACG','complex' | 0.007 | missense_variant c.357_363delAGATGCTinsGGAAACG p.AspAla120GluThr | |
| '1420295','AATAGGT:CATTGGC','complex' | 0.006 | synonymous_variant c.276_282delAATAGGTinsCATTGGC p.95 | |
| '1587885','T:G','snp' | 0.006 | synonymous_variant c.414A>C p.Leu138Leu | |
| '967881','C:T','snp' | 0.005 | missense_variant c.242C>T p.Ser81Leu | gyrA |
| '935653','A:G','snp' | 0.005 | synonymous_variant c.3033A>G p.Glu1011Glu | posA |

**Table 43 – continued from previous page**

| Feature Name | Gini Importance | Mutation Annotation | Gene |
|---|---|---|---|
| '672496','T:A','snp' | 0.005 | | |

#### Acinetobacter baumannii - trimethoprim_sulfamethoxazole

| Bacteria | Antibiotic | F1 Score | MCC |
|---|---|---|---|
| Acinetobacter baumannii | trimethoprim_sulfamethoxazole | 0.946 | 0.521 |

Table 44: Feature Importance for Acinetobacter baumannii - trimethoprim_sulfamethoxazole

| Feature Name | Gini Importance | Mutation Annotation | Gene |
|---|---|---|---|
| '3012426','T:C','snp' | 0.008 | synonymous_variant c.931T>C p.Leu311Leu | |
| '2410408','TA: CG','complex' | 0.008 | missense_variant c.584_585delTAinsCG p.Val195Ala | |
| 051L7659_IOEMPANA_02909_dihydropteroate_synthase | 0.006 | | |
| 0129PRKG_JOKDIMCI_02673_hypothetical_protein | 0.006 | | |
| '2399725','G:A','snp' | 0.006 | synonymous_variant c.486C>T p.Ser162Ser | dcaP |
| 0397U9GK_FKALAFKA_03899_Phosphoglucosamine_mutase | 0.005 | | |
| '1998430','CAAAATA: TAAGCTT','complex' | 0.004 | missense_variant c.669_675delCAAAATAinsTAAGCTT p.Ile225Leu | |
| '2383982','G:A','snp' | 0.004 | synonymous_variant c.159G>A p.Lys53Lys | |
| JZ8YMCRP_KCFCEHDF_00338_hypothetical_protein | 0.003 | | |
| '1692893','G:A','snp' | 0.003 | synonymous_variant c.186C>T p.Ala62Ala | argF |

#### Acinetobacter baumannii - imipenem

| Bacteria | Antibiotic | F1 Score | MCC |
|---|---|---|---|
| Acinetobacter baumannii | imipenem | 0.900 | 0.810 |

Table 45: Feature Importance for Acinetobacter baumannii - imipenem

| Feature Name | Gini Importance | Mutation Annotation | Gene |
|---|---|---|---|
| '3024605','C:T','snp' | 0.005 | synonymous_variant c.540G>A p.Leu180Leu | |
| '3024752','A:G','snp' | 0.005 | synonymous_variant c.393T>C p.Thr131Thr | |
| '3024914','A:G','snp' | 0.005 | synonymous_variant c.231T>C p.Ala77Ala | |
| '3017375','G:A','snp' | 0.005 | synonymous_variant c.876C>T p.Ile292Ile | |
| '3023876','T:C','snp' | 0.005 | synonymous_variant c.114A>G p.Gln38Gln | |
| '3024299','G:A','snp' | 0.004 | synonymous_variant c.846C>T p.Arg282Arg | |
| '2868127','GACCA: AACCG','complex' | 0.004 | missense_variant c.234_238delTGGTCinsCGGTT p.Pro80Ser | |
| '3015643','GTTC: ATTA','complex' | 0.003 | | |
| '3022402','G:T','snp' | 0.003 | synonymous_variant c.1588C>A p.Arg530Arg | |
| '3022424','C:T','snp' | 0.003 | synonymous_variant c.1566G>A p.Gly522Gly | |

#### Acinetobacter baumannii - amikacin

| Bacteria | Antibiotic | F1 Score | MCC |
|---|---|---|---|
| Acinetobacter baumannii | amikacin | 0.913 | 0.802 |

Table 46: Feature Importance for Acinetobacter baumannii - amikacin

| Feature Name | Gini Importance | Mutation Annotation | Gene |
|---|---|---|---|
| 0208T4X2_JJPNOEOJ_00715_Aminoglycoside_3__phosphotransferase | 0.010 | | |
| '1328622','AGAA: TGCT','complex' | 0.004 | missense_variant c.720_723delTTCTinsAGCA p.Ser241Ala | adeF |
| '2357474','G:T','snp' | 0.003 | synonymous_variant c.1155C>A p.Gly385Gly | |
| 02K1STUU_ECCICHIJ_01947_hypothetical_protein | 0.003 | | |
| '1859163','G:A','snp' | 0.003 | synonymous_variant c.126C>T p.Ser42Ser | |

**Table 46 – continued from previous page**

| Feature Name | Gini Importance | Mutation Annotation | Gene |
|---|---|---|---|
| 02K1STUU_ECCICHIJ_01907_hypothetical_protein | 0.003 | | |
| '2645505','TT:CG','complex' | 0.003 | missense_variant c.9_10delTTinsCG p.Leu4Val | |
| '2829774','TA:T','del' | 0.003 | | |
| '1899189','T:A','snp' | 0.003 | missense_variant c.2266T>A p.Ser756Thr | |
| 02K1STUU_ECCICHIJ_01938_ATP_dependent_RecD_like_DNA_helicase | 0.003 | | |

#### Acinetobacter baumannii - ceftazidime

| Bacteria | Antibiotic | F1 Score | MCC |
|---|---|---|---|
| Acinetobacter baumannii | ceftazidime | 0.968 | 0.801 |

Table 47: Feature Importance for Acinetobacter baumannii - ceftazidime

| Feature Name | Gini Importance | Mutation Annotation | Gene |
|---|---|---|---|
| '3557211','CA:AG','complex' | 0.014 | | |
| '2941422','C:T','snp' | 0.013 | splice_region_variant&stop_retained_variant c.614G>A p.Ter205Ter | |
| '2189486','T:C','snp' | 0.012 | synonymous_variant c.450A>G p.Glu150Glu | |
| '1905229','A:C','snp' | 0.011 | synonymous_variant c.303A>C p.Ser101Ser | |
| '820158','T:C','snp' | 0.011 | synonymous_variant c.219A>G p.Val73Val | |
| '722619','A:T','snp' | 0.011 | synonymous_variant c.498T>A p.Ile166Ile | |
| '3663821','C:T','snp' | 0.010 | synonymous_variant c.2496G>A p.Gln832Gln | rpoC |
| '2686451','G:A','snp' | 0.010 | synonymous_variant c.315C>T p.Val105Val | |
| '1632948','T:C','snp' | 0.009 | synonymous_variant c.90T>C p.Gly30Gly | |
| '2092345','C:T','snp' | 0.007 | synonymous_variant c.48C>T p.Ser16Ser | |

#### Acinetobacter baumannii - tobramycin

| Bacteria | Antibiotic | F1 Score | MCC |
|---|---|---|---|
| Acinetobacter baumannii | tobramycin | 0.924 | 0.749 |

Table 48: Feature Importance for Acinetobacter baumannii - tobramycin

| Feature Name | Gini Importance | Mutation Annotation | Gene |
|---|---|---|---|
| '3781217','A:C','snp' | 0.005 | synonymous_variant c.372A>C p.Gly124Gly | |
| '3748081','C:G','snp' | 0.005 | synonymous_variant c.87G>C p.Thr29Thr | |
| '3829856','A:G','snp' | 0.005 | synonymous_variant c.408T>C p.Asn136Asn | abaR |
| '3965802','C:A','snp' | 0.004 | synonymous_variant c.408G>T p.Leu136Leu | |
| 02K1STUU_ECCICHIJ_02422_hypothetical_protein | 0.004 | | |
| '2829803','C:T','snp' | 0.004 | | |
| '3814285','G:A','snp' | 0.003 | synonymous_variant c.1722C>T p.Thr574Thr | |
| '3814246','G:A','snp' | 0.003 | synonymous_variant c.1761C>T p.Gly587Gly | |
| '24025','A:C','snp' | 0.003 | synonymous_variant c.390A>C p.Pro130Pro | |
| '2831001','T:C','snp' | 0.003 | synonymous_variant c.1083T>C p.Ile361Ile | |

#### Acinetobacter baumannii - levofloxacin

| Bacteria | Antibiotic | F1 Score | MCC |
|---|---|---|---|
| Acinetobacter baumannii | levofloxacin | 0.956 | 0.782 |

Table 49: Feature Importance for Acinetobacter baumannii - levofloxacin

| Feature Name | Gini Importance | Mutation Annotation | Gene |
|---|---|---|---|
| '2610939','C:A','snp' | 0.013 | missense_variant c.289C>A p.Leu97Met | |
| '2937464','C:A','snp' | 0.012 | missense_variant c.711C>A p.Phe237Leu | |
| '3293834','A:T','snp' | 0.010 | synonymous_variant c.102T>A p.Thr34Thr | hisG |
| '1144491','C:A','snp' | 0.010 | synonymous_variant c.33C>A p.Val11Val | surE |
| '3321139','G:A','snp' | 0.008 | synonymous_variant c.267C>T p.Tyr89Tyr | |

**Table 49 – continued from previous page**

| Feature Name | Gini Importance | Mutation Annotation | Gene |
|---|---|---|---|
| '1115587','G:A','snp' | 0.007 | synonymous_variant c.240G>A p.Thr80Thr | |
| '1362346','GGCC:AGCT','complex' | 0.006 | synonymous_variant c.765_768delGGCCinsAGCT p.257 | |
| '1420307','A:G','snp' | 0.006 | synonymous_variant c.288A>G p.Leu96Leu | |
| '3557211','CA:AG','complex' | 0.006 | | |
| '2882172','CTAT:ATA','complex' | 0.006 | | |

#### Acinetobacter baumannii - carbapenem

| Bacteria | Antibiotic | F1 Score | MCC |
|---|---|---|---|
| Acinetobacter baumannii | carbapenem | 0.939 | 0.873 |

Table 50: Feature Importance for Acinetobacter baumannii - carbapenem

| Feature Name | Gini Importance | Mutation Annotation | Gene |
|---|---|---|---|
| '1890237','TGCT:CGCC','complex' | 0.013 | synonymous_variant c.216_219delTGCTinsCGCC p.74 | |
| '3358151','A:T','snp' | 0.013 | synonymous_variant c.300A>T p.Pro100Pro | |
| '3182471','T:TA','ins' | 0.008 | | |
| '1585254','T:G','snp' | 0.008 | synonymous_variant c.1212T>G p.Ala404Ala | |
| '3608561','ACTG:GCTA','complex' | 0.007 | synonymous_variant c.903_906delACTGinsGCTA p.303 | |
| '3018047','C:T','snp' | 0.007 | synonymous_variant c.204G>A p.Val68Val | |
| '3023099','A:G','snp' | 0.007 | synonymous_variant c.891T>C p.Phe297Phe | |
| '525190','G:A','snp' | 0.007 | synonymous_variant c.291C>T p.Asn97Asn | |
| '1151685','T:C','snp' | 0.007 | synonymous_variant c.1641T>C p.Gly547Gly | |
| '664808','G:A','snp' | 0.007 | synonymous_variant c.2280C>T p.Thr760Thr | |

#### Acinetobacter baumannii - ampicillin_sulbactam

| Bacteria | Antibiotic | F1 Score | MCC |
|---|---|---|---|
| Acinetobacter baumannii | ampicillin_sulbactam | 0.918 | 0.836 |

Table 51: Feature Importance for Acinetobacter baumannii - ampicillin_sulbactam

| Feature Name | Gini Importance | Mutation Annotation | Gene |
|---|---|---|---|
| '1223356','T:G','snp' | 0.009 | synonymous_variant c.924T>G p.Gly308Gly | xdhA |
| '1561705','T:G','snp' | 0.009 | missense_variant c.124T>G p.Ser42Ala | |
| '1016484','A:G','snp' | 0.006 | synonymous_variant c.2997A>G p.Gln999Gln | purL |
| '556087','G:A','snp' | 0.005 | synonymous_variant c.219G>A p.Glu73Glu | |
| '3567042','C:T','snp' | 0.005 | | |
| '3139042','G:GT','ins' | 0.005 | | |
| '2925062','A:G','snp' | 0.005 | synonymous_variant c.1128T>C p.Gly376Gly | |
| 0129PRKG_JOKDIMCI_03749_phage_tail_protein | 0.005 | | |
| '864497','A:T','snp' | 0.005 | synonymous_variant c.855A>T p.Ser285Ser | |
| '2521714','TAATTC:CAACTG','complex' | 0.005 | missense_variant c.58_63delGAATTAinsCAGTTG p.Glu20Gln | |

#### Acinetobacter baumannii - tetracycline

| Bacteria | Antibiotic | F1 Score | MCC |
|---|---|---|---|
| Acinetobacter baumannii | tetracycline | 0.969 | 0.863 |

Table 52: Feature Importance for Acinetobacter baumannii - tetracycline

| Feature Name | Gini Importance | Mutation Annotation | Gene |
|---|---|---|---|
| '2829878','G:A','snp' | 0.016 | | |
| '3721169','T:C','snp' | 0.011 | synonymous_variant c.333A>G p.Leu111Leu | lptG |
| '3352194','G:A','snp' | 0.011 | synonymous_variant c.900C>T p.Phe300Phe | cysM |
| '1632631','A:G','snp' | 0.010 | synonymous_variant c.1239A>G p.Ala413Ala | |
| '2338993','C:A','snp' | 0.007 | synonymous_variant c.747C>A p.Thr249Thr | |

**Table 52 – continued from previous page**

| Feature Name | Gini Importance | Mutation Annotation | Gene |
|---|---|---|---|
| '1133755','T:C','snp' | 0.007 | synonymous_variant c.396T>C p.Val132Val | sohB |
| '2347300','A:T','snp' | 0.007 | synonymous_variant c.1833A>T p.Ala611Ala | |
| '775583','TAGC: CAGT','complex' | 0.006 | synonymous_variant c.234_237delTAGCinsCAGT p.80 | |
| '2347324','TGTG: CGTC','complex' | 0.006 | synonymous_variant c.1857_1860delTGTGinsCGTC p.621 | |
| '2339352','C:T','snp' | 0.006 | synonymous_variant c.81C>T p.His27His | yfcF |

#### Acinetobacter baumannii - meropenem

| Bacteria | Antibiotic | F1 Score | MCC |
|---|---|---|---|
| Acinetobacter baumannii | meropenem | 0.892 | 0.701 |

Table 53: Feature Importance for Acinetobacter baumannii - meropenem

| Feature Name | Gini Importance | Mutation Annotation | Gene |
|---|---|---|---|
| 2P6FYN27_OCDCJDIH_00132_ATP_binding_protein | 0.006 | | |
| '3016163','CCCAGAAGTG:TCCTGAAATA','complex' | 0.005 | missense_variant c.213_222delCCCAGAAGTGinsTCCTGAAATA p.Val74Ile | |
| 1GEMOD5L_DNOOALJC_02653_IS5_family_transposase_ISEc35 | 0.005 | | |
| 02K1STUU_ECCICHIJ_02735_hypothetical_protein | 0.005 | | |
| '3022541','A:G','snp' | 0.005 | synonymous_variant c.1449T>C p.Ile483Ile | |
| 02K1STUU_ECCICHIJ_03425_replication_initiation_protein_RepM | 0.004 | | |
| 02K1STUU_ECCICHIJ_02709_hypothetical_protein | 0.004 | | |
| '1471918','T:C','snp' | 0.004 | missense_variant c.355T>C p.Tyr119His | |
| 02K1STUU_ECCICHIJ_02741_hypothetical_protein | 0.003 | | |

**Table 53 – continued from previous page**

| Feature Name | Gini Importance | Mutation Annotation | Gene |
|---|---|---|---|
| 02K1STUU_ECCICHIJ_02722_hypothetical_protein | 0.003 | | |

#### Klebsiella pneumoniae - ceftazidime

| Bacteria | Antibiotic | F1 Score | MCC |
|---|---|---|---|
| Klebsiella pneumoniae | ceftazidime | 0.969 | 0.642 |

Table 54: Feature Importance for Klebsiella pneumoniae - ceftazidime

| Feature Name | Gini Importance | Mutation Annotation | Gene |
|---|---|---|---|
| '1705353','A:G','snp' | 0.007 | synonymous_variant c.643T>C p.Leu215Leu | |
| '3015951','T:A','snp' | 0.007 | missense_variant c.346A>T p.Thr116Ser | |
| '5296733','TAAACGC:CAGACGG','complex' | 0.006 | synonymous_variant c.216_222delGCGTTTAinsCCGTCTG p.75 | |
| '2711665','T:C','snp' | 0.005 | | |
| '3031310','GTAT:ATAA','complex' | 0.005 | synonymous_variant c.2652_2655delATACinsTTAT p.886 | |
| '3041653','T:C','snp' | 0.005 | | |
| '2816604','TT:CA','complex' | 0.004 | missense_variant c.17_18delTTinsCA p.Leu6Pro | |
| '3387961','G:A','snp' | 0.004 | synonymous_variant c.376C>T p.Leu126Leu | |
| '3041785','T:C','snp' | 0.004 | | |
| '1537348','A:G','snp' | 0.004 | synonymous_variant c.1365T>C p.Asp455Asp | |

#### Klebsiella pneumoniae - levofloxacin

| Bacteria | Antibiotic | F1 Score | MCC |
|---|---|---|---|
| Klebsiella pneumoniae | levofloxacin | 0.961 | 0.846 |

Table 55: Feature Importance for Klebsiella pneumoniae - levofloxacin

| Feature Name | Gini Importance | Mutation Annotation | Gene |
|---|---|---|---|
| E07YHER4_BCOPJOCI_04862_Outer_membrane_protein_YedS | 0.014 | | |

**Table 55 – continued from previous page**

| Feature Name | Gini Importance | Mutation Annotation | Gene |
|---|---|---|---|
| '3015940','A:G','snp' | 0.012 | synonymous_variant c.357T>C p.Gly119Gly | |
| '3268347','C:T','snp' | 0.011 | synonymous_variant c.69G>A p.Leu23Leu | |
| '3303550','G:A','snp' | 0.010 | synonymous_variant c.391C>T p.Leu131Leu | |
| '3388862','C:T','snp' | 0.009 | synonymous_variant c.174C>T p.Arg58Arg | |
| TTNLMLPO_MGMLNCPH_03513_putative_oxidoreductase | 0.008 | | |
| '306144','T:C','snp' | 0.008 | synonymous_variant c.357T>C p.Ser119Ser | |
| 01UJQEZO_FDFJBMGD_01556_hypothetical_protein | 0.007 | | |
| 01UJQEZO_FDFJBMGD_01555_hypothetical_protein | 0.006 | | |
| '3388631','G:A','snp' | 0.006 | | |

#### Klebsiella pneumoniae - tetracycline

| Bacteria | Antibiotic | F1 Score | MCC |
|---|---|---|---|
| Klebsiella pneumoniae | tetracycline | 0.747 | 0.503 |

Table 56: Feature Importance for Klebsiella pneumoniae - tetracycline

| Feature Name | Gini Importance | Mutation Annotation | Gene |
|---|---|---|---|
| IBCMBDAS_JPFBIIEK_05357_tetracycline_resistance_protein | 0.006 | | |
| 001QJFFS_HPLJBCIE_00040_tetracycline_resistance_protein | 0.006 | | |
| 03VPBTRU_MPBKBMBA_05289_alcohol_dehydrogenase_class_III | 0.004 | | |
| 001QJFFS_HPLJBCIE_00039_tetracycline_repressor_protein_class_G | 0.004 | | |
| 001QJFFS_HPLJBCIE_00041_putative_transmembrane_protein | 0.003 | | |

**Table 56 – continued from previous page**

| Feature Name | Gini Importance | Mutation Annotation | Gene |
|---|---|---|---|
| '133690','G:A','snp' | 0.003 | synonymous_variant c.1290G>A p.Ala430Ala | |
| '3061317','T:C','snp' | 0.003 | synonymous_variant c.993T>C p.Gly331Gly | |
| '5178548','A:G','snp' | 0.003 | synonymous_variant c.291A>G p.Leu97Leu | |
| '1120938','T:C','snp' | 0.003 | synonymous_variant c.172T>C p.Leu58Leu | |
| '677156','C:T','snp' | 0.003 | synonymous_variant c.192G>A p.Ala64Ala | |

#### Klebsiella pneumoniae - cefazolin

| Bacteria | Antibiotic | F1 Score | MCC |
|---|---|---|---|
| Klebsiella pneumoniae | cefazolin | 0.979 | 0.784 |

Table 57: Feature Importance for Klebsiella pneumoniae - cefazolin

| Feature Name | Gini Importance | Mutation Annotation | Gene |
|---|---|---|---|
| '2943262','C:T','snp' | 0.013 | synonymous_variant c.978C>T p.Gly326Gly | |
| '2121329','A:G','snp' | 0.009 | synonymous_variant c.192A>G p.Val64Val | |
| '3738483','AT: GA','complex' | 0.008 | missense_variant c.247_248delATinsTC p.Ile83Ser | |
| '3041653','T:C','snp' | 0.008 | | |
| '1543581','T:C','snp' | 0.008 | missense_variant c.211A>G p.Thr71Ala | |
| '2165083','A:C','snp' | 0.007 | synonymous_variant c.180T>G p.Pro60Pro | |
| '108490','A:G','snp' | 0.005 | synonymous_variant c.447T>C p.Ala149Ala | |
| '1657694','A:C','snp' | 0.005 | synonymous_variant c.321T>G p.Arg107Arg | |
| A2AJPJ11_HMDMFNFI_09856_Pentapeptide_repeat_protein | 0.005 | | |
| '3231302','G:A','snp' | 0.005 | synonymous_variant c.153C>T p.Phe51Phe | |

#### Klebsiella pneumoniae - gentamicin

| Bacteria | Antibiotic | F1 Score | MCC |
|---|---|---|---|
| Klebsiella pneumoniae | gentamicin | 0.880 | 0.805 |

Table 58: Feature Importance for Klebsiella pneumoniae - gentamicin

| Feature Name | Gini Importance | Mutation Annotation | Gene |
|---|---|---|---|
| '44569','G:A','snp' | 0.015 | missense_variant c.493G>A p.Asp165Asn | |
| '44914','A:G','snp' | 0.011 | missense_variant c.52A>G p.Lys18Glu | |
| '44517','C:G','snp' | 0.010 | synonymous_variant c.441C>G p.Ala147Ala | |
| '44658','T:C','snp' | 0.009 | synonymous_variant c.582T>C p.Val194Val | |
| '44439','T:C','snp' | 0.007 | synonymous_variant c.363T>C p.Val121Val | |
| '44418','A:G','snp' | 0.007 | synonymous_variant c.342A>G p.Gly114Gly | |
| '44598','GGAT: AGAA','complex' | 0.006 | missense_variant c.522_525delGGATinsAGAA p.Asp175Glu | |
| '44367','G:A','snp' | 0.006 | synonymous_variant c.291G>A p.Leu97Leu | |
| '44784','T:C','snp' | 0.005 | synonymous_variant c.708T>C p.His236His | |
| '44844','A:C','snp' | 0.004 | synonymous_variant c.768A>C p.Ser256Ser | |

#### Klebsiella pneumoniae - trimethoprim_sulfamethoxazole

| Bacteria | Antibiotic | F1 Score | MCC |
|---|---|---|---|
| Klebsiella pneumoniae | trimethoprim_sulfamethoxazole | 0.929 | 0.745 |

Table 59: Feature Importance for Klebsiella pneumoniae - trimethoprim_sulfamethoxazole

| Feature Name | Gini Importance | Mutation Annotation | Gene |
|---|---|---|---|
| IUD3RWWZ_JDJMNOMM_05050_putative_transposase | 0.009 | | |
| 06S1BQ7K_OHPILBJP_05126_hypothetical_protein | 0.008 | | |
| H2TWJMMS_GPKPNAOG_05017_class_I_integron_integrase | 0.008 | | |
| '53703','A:C','snp' | 0.004 | | |
| 01UJQEZO_FDFJBMGD_05044_hypothetical_protein | 0.004 | | |

**Table 59 – continued from previous page**

| Feature Name | Gini Importance | Mutation Annotation | Gene |
|---|---|---|---|
| 45I3LUB9_ACHJPDAC_03955_Antirestriction_protein_KlcA | 0.004 | | |
| '42183','C:T','snp' | 0.004 | synonymous_variant c.90C>T p.Ser30Ser | |
| JN848DOI_JJHFIICA_05300_Tn4653_resolvase | 0.004 | | |
| 06S1BQ7K_OHPILBJP_03830_dihydrofolate_reductase | 0.003 | | |
| J7JXZC4W_FEFDNFBP_05405_Spermidine_export_protein_MdtJ | 0.003 | | |

#### Klebsiella pneumoniae - ceftriaxone

| Bacteria | Antibiotic | F1 Score | MCC |
|---|---|---|---|
| Klebsiella pneumoniae | ceftriaxone | 0.979 | 0.772 |

Table 60: Feature Importance for Klebsiella pneumoniae - ceftriaxone

| Feature Name | Gini Importance | Mutation Annotation | Gene |
|---|---|---|---|
| '3103680','G:A','snp' | 0.009 | synonymous_variant c.1755C>T p.Ala585Ala | |
| '818030','T:C','snp' | 0.007 | synonymous_variant c.420A>G p.Ala140Ala | |
| '1190804','C:G','snp' | 0.007 | synonymous_variant c.588G>C p.Ala196Ala | |
| 006F75LC_HJOODNNJ_05248_hypothetical_protein | 0.007 | | |
| '3867336','G:T','snp' | 0.006 | synonymous_variant c.324G>T p.Ala108Ala | |
| '3015951','T:A','snp' | 0.006 | missense_variant c.346A>T p.Thr116Ser | |
| '4779790','TGCC:CGCT','complex' | 0.006 | synonymous_variant c.3171_3174delTGCCinsCGCT p.1059 | |
| '4591623','A:C','snp' | 0.006 | missense_variant c.239T>G p.Ile80Ser | |
| 276QJ4DR_LKONKJAA_00765_hypothetical_protein | 0.005 | | |
| '2821033','G:A','snp' | 0.005 | missense_variant c.773C>T p.Pro258Leu | |

#### Klebsiella pneumoniae - amikacin

| Bacteria | Antibiotic | F1 Score | MCC |
|---|---|---|---|
| Klebsiella pneumoniae | amikacin | 0.733 | 0.702 |

Table 61: Feature Importance for Klebsiella pneumoniae - amikacin

| Feature Name | Gini Importance | Mutation Annotation | Gene |
|---|---|---|---|
| '4500270','G:A','snp' | 0.008 | synonymous_variant c.1263C>T p.Ile421Ile | |
| '3721530','C:T','snp' | 0.008 | synonymous_variant c.1023C>T p.Thr341Thr | |
| '4103518','TACT: CACC','complex' | 0.007 | synonymous_variant c.243_246delTACTinsCACC p.83 | |
| '2830835','GACG: TACA','complex' | 0.007 | synonymous_variant c.684_687delCGTCinsTGTA p.230 | |
| '2872107','G:A','snp' | 0.005 | synonymous_variant c.399C>T p.Arg133Arg | |
| '3303065','G:A','snp' | 0.005 | synonymous_variant c.876C>T p.Arg292Arg | |
| '3414426','TGTC: GGTA','complex' | 0.004 | synonymous_variant c.363_366delTGTCinsGGTA p.123 | |
| '539365','T:C','snp' | 0.004 | synonymous_variant c.156A>G p.Glu52Glu | |
| '4849907','C:T','snp' | 0.004 | synonymous_variant c.165G>A p.Glu55Glu | |
| '4655126','A:G','snp' | 0.004 | missense_variant c.919A>G p.Thr307Ala | |

#### Klebsiella pneumoniae - aztreonam

| Bacteria | Antibiotic | F1 Score | MCC |
|---|---|---|---|
| Klebsiella pneumoniae | aztreonam | 0.936 | 0.461 |

Table 62: Feature Importance for Klebsiella pneumoniae - aztreonam

| Feature Name | Gini Importance | Mutation Annotation | Gene |
|---|---|---|---|
| 006F75LC_HJOODNNJ_05247_beta_lactamase_CTX_M_14 | 0.006 | | |
| '2900204','C:T','snp' | 0.005 | missense_variant c.229G>A p.Ala77Thr | |
| '3287971','G:A','snp' | 0.004 | missense_variant c.403C>T p.Leu135Phe | |

**Table 62 – continued from previous page**

| Feature Name | Gini Importance | Mutation Annotation | Gene |
|---|---|---|---|
| '5237907','T:C','snp' | 0.003 | synonymous_variant c.993A>G p.Thr331Thr | |
| '2639591','CGAA: AGAC','complex' | 0.003 | missense_variant c.594_597delTTCGinsGTCT p.Phe198Leu | |
| '3738483','AT: GA','complex' | 0.003 | missense_variant c.247_248delATinsTC p.Ile83Ser | |
| '2717127','CCGCCCCC: GCGCTCA','complex' | 0.003 | | |
| '2487924','A:G','snp' | 0.003 | synonymous_variant c.610T>C p.Leu204Leu | |
| '2933473','A:T','snp' | 0.003 | synonymous_variant c.30A>T p.Leu10Leu | |
| 05GCBKNR_FPLHKBIC_00018_hypothetical_protein | 0.003 | | |

#### Klebsiella pneumoniae - piperacillin_tazobactam

| Bacteria | Antibiotic | F1 Score | MCC |
|---|---|---|---|
| Klebsiella pneumoniae | piperacillin_tazobactam | 0.897 | 0.669 |

Table 63: Feature Importance for Klebsiella pneumoniae - piperacillin_tazobactam

| Feature Name | Gini Importance | Mutation Annotation | Gene |
|---|---|---|---|
| '1685878','T:C','snp' | 0.006 | missense_variant c.1009A>G p.Thr337Ala | |
| '2122390','A:G','snp' | 0.006 | missense_variant c.712T>C p.Ser238Pro | |
| 01UJQEZO_FDFJBMGD_04983_IS1182_family_transposase_ISKpn6 | 0.006 | | |
| '2253433','ATT: CTC','complex' | 0.006 | missense_variant c.256_258delATTinsCTC p.Ile86Leu | |
| '5205584','T:C','snp' | 0.006 | missense_variant c.508A>G p.Ile170Val | |
| '1423061','A:G','snp' | 0.004 | synonymous_variant c.774A>G p.Leu258Leu | |
| '3867336','G:T','snp' | 0.004 | synonymous_variant c.324G>T p.Ala108Ala | |
| '3404411','C:A','snp' | 0.004 | missense_variant c.644G>T p.Cys215Phe | |
| '4254587','A:G','snp' | 0.004 | synonymous_variant c.210T>C p.Ser70Ser | |
| '306144','T:C','snp' | 0.004 | synonymous_variant c.357T>C p.Ser119Ser | |

#### Klebsiella pneumoniae - meropenem

| Bacteria | Antibiotic | F1 Score | MCC |
|---|---|---|---|
| Klebsiella pneumoniae | meropenem | 0.881 | 0.820 |

Table 64: Feature Importance for Klebsiella pneumoniae - meropenem

| Feature Name | Gini Importance | Mutation Annotation | Gene |
|---|---|---|---|
| 01UJQEZO_FDFJBMGD_04982_Class_A_Carbapenemase_Kpc_2 | 0.014 | | |
| T9ACDQPA_GKLPDKGO_04755_hypothetical_protein | 0.009 | | |
| PK5LTS6E_IMIKPLMD_05005_RNA_chaperone_ProQ | 0.009 | | |
| Q116SWRB_MHIIGNEK_05725_Tn3_family_transposase_ISPsy42 | 0.006 | | |
| 01UJQEZO_FDFJBMGD_04980_hypothetical_protein | 0.006 | | |
| '1758220','T:C','snp' | 0.005 | synonymous_variant c.471T>C p.Pro157Pro | |
| '813092','C:G','snp' | 0.005 | missense_variant c.2750G>C p.Gly917Ala | |
| '1158750','A:G','snp' | 0.005 | synonymous_variant c.102A>G p.Leu34Leu | |
| 01UJQEZO_FDFJBMGD_04978_Tyrosine_recombinase_XerC | 0.005 | | |
| YLAVYB8I_FLGBLIPD_00987_type_F_conjugative_transfer_system_pilin_acetylase_TraX | 0.004 | | |

#### Klebsiella pneumoniae - cefepime

| Bacteria | Antibiotic | F1 Score | MCC |
|---|---|---|---|
| Klebsiella pneumoniae | cefepime | 0.843 | 0.588 |

Table 65: Feature Importance for Klebsiella pneumoniae - cefepime

| Feature Name | Gini Importance | Mutation Annotation | Gene |
|---|---|---|---|
| L5EHL7HU_CDMPAKHP_05726_Aminoglycoside_N_6___acetyltransferase_type_1 | 0.005 | | |
| 06S1BQ7K_OHPILBJP_00729_hypothetical_protein | 0.004 | | |
| 04SIPUD6_CJMCGLKO_01883_hypothetical_protein | 0.004 | | |
| NBB8QBUE_DCHKCAKD_05500_transposase_for_transposon | 0.004 | | |
| '1910554','A:ACCGATTATTGCCTGGCTATCG','ins' | 0.004 | disruptive_inframe_insertion c.903_923dupTGCCTGGCTATCGCCGATTAT p.Ile308_Ala309insAlaTrpLeuSerProIleIle | |
| 006F75LC_HJOODNNJ_05247_beta_lactamase_CTX_M_14 | 0.004 | | |
| '2890135','C:A','snp' | 0.004 | synonymous_variant c.147G>T p.Ser49Ser | |
| '3376493','CTCA:TTCG','complex' | 0.003 | synonymous_variant c.1731_1734delTGAGinsCGAA p.579 | |
| 006F75LC_HJOODNNJ_05248_hypothetical_protein | 0.003 | | |
| 001QJFFS_HPLJBCIE_04861_extended_spectrum_beta_lactamase_aminoglycoside_modifying_enzyme_fusion_protein | 0.003 | | |

#### Klebsiella pneumoniae - imipenem

| Bacteria | Antibiotic | F1 Score | MCC |
|---|---|---|---|
| Klebsiella pneumoniae | imipenem | 0.873 | 0.794 |

Table 66: Feature Importance for Klebsiella pneumoniae - imipenem

| Feature Name | Gini Importance | Mutation Annotation | Gene |
|---|---|---|---|
| 01UJQEZO_FDFJBMGD_04983_IS1182_family_transposase_ISKpn6 | 0.015 | | |
| 01UJQEZO_FDFJBMGD_04982_Class_A_Carbapenemase_Kpc_2 | 0.012 | | |
| 01UJQEZO_FDFJBMGD_04981_IS21_family_transposase_ISKpn7 | 0.012 | | |
| LH50ZBDW_OOEHFJIF_05562_ISAs1_family_transposase_ISKpn31 | 0.008 | | |
| T9ACDQPA_GKLPDKGO_04756_hypothetical_protein | 0.006 | | |
| 01UJQEZO_FDFJBMGD_04985_hypothetical_protein | 0.005 | | |
| '511896','A:G','snp' | 0.005 | synonymous_variant c.456T>C p.Ile152Ile | |
| '1760202','T:G','snp' | 0.005 | synonymous_variant c.561T>G p.Pro187Pro | |
| '2028969','A:G','snp' | 0.005 | | |
| '101316','G:A','snp' | 0.004 | | |

#### Klebsiella pneumoniae - cefoxitin

| Bacteria | Antibiotic | F1 Score | MCC |
|---|---|---|---|
| Klebsiella pneumoniae | cefoxitin | 0.861 | 0.684 |

Table 67: Feature Importance for Klebsiella pneumoniae - cefoxitin

| Feature Name | Gini Importance | Mutation Annotation | Gene |
|---|---|---|---|
| '5068070','A:G','snp' | 0.007 | synonymous_variant c.1188T>C p.Thr396Thr | |
| '5008633','A:G','snp' | 0.007 | synonymous_variant c.696A>G p.Gly232Gly | |
| '4982418','A:G','snp' | 0.007 | | |
| 01UJQEZO_FDFJBMGD_04983_IS1182_family_transposase_ISKpn6 | 0.007 | | |
| 01UJQEZO_FDFJBMGD_04982_Class_A_Carbapenemase_Kpc_2 | 0.006 | | |

**Table 67 – continued from previous page**

| Feature Name | Gini Importance | Mutation Annotation | Gene |
|---|---|---|---|
| '1774612','T:C','snp' | 0.006 | synonymous_variant c.276T>C p.Phe92Phe | |
| '3941728','T:C','snp' | 0.005 | synonymous_variant c.1257T>C p.Thr419Thr | |
| 01UJQEZO_FDFJBMGD_04980_hypothetical_protein | 0.005 | | |
| '406185','A:G','snp' | 0.004 | synonymous_variant c.1344T>C p.Tyr448Tyr | |
| '4789245','G:A','snp' | 0.004 | | |

#### Klebsiella pneumoniae - ciprofloxacin

| Bacteria | Antibiotic | F1 Score | MCC |
|---|---|---|---|
| Klebsiella pneumoniae | ciprofloxacin | 0.965 | 0.818 |

Table 68: Feature Importance for Klebsiella pneumoniae - ciprofloxacin

| Feature Name | Gini Importance | Mutation Annotation | Gene |
|---|---|---|---|
| '2928634','A:T','snp' | 0.013 | | |
| '2946958','G:T','snp' | 0.012 | synonymous_variant c.24G>T p.Ala8Ala | |
| '2711635','CTGA:TTGT','complex' | 0.009 | | |
| '3741209','T:C','snp' | 0.007 | synonymous_variant c.1260T>C p.Cys420Cys | |
| '2660159','A:G','snp' | 0.007 | synonymous_variant c.930A>G p.Ala310Ala | |
| '3634483','C:T','snp' | 0.006 | synonymous_variant c.144C>T p.Val48Val | |
| '2943262','C:T','snp' | 0.006 | synonymous_variant c.978C>T p.Gly326Gly | |
| '520100','TC:T','del' | 0.006 | | |
| '30574','T:G','snp' | 0.006 | synonymous_variant c.360T>G p.Leu120Leu | |
| '1397852','G:C','snp' | 0.006 | synonymous_variant c.471C>G p.Ala157Ala | |

#### Klebsiella pneumoniae - ertapenem

| Bacteria | Antibiotic | F1 Score | MCC |
|---|---|---|---|
| Klebsiella pneumoniae | ertapenem | 0.913 | 0.684 |

Table 69: Feature Importance for Klebsiella pneumoniae - ertapenem

| Feature Name | Gini Importance | Mutation Annotation | Gene |
|---|---|---|---|
| '2880928','C:G','snp' | 0.010 | synonymous_variant c.582C>G p.Ser194Ser | |
| '3274905','TCGAC: ACGAT','complex' | 0.007 | missense_variant c.216_220delGTCGAinsATCGT p.Thr74Ser | |
| '2720229','TTCC: CTCA','complex' | 0.006 | missense_variant c.927_930delGGAAinsTGAG p.Glu309Asp | |
| '3528765','A:G','snp' | 0.006 | synonymous_variant c.138A>G p.Glu46Glu | |
| 05GCBKNR_FPLHKBIC_01266_hypothetical_protein | 0.005 | | |
| '2612321','G:C','snp' | 0.004 | synonymous_variant c.351C>G p.Gly117Gly | |
| '3528594','C:T','snp' | 0.004 | | |
| '3646166','C:T','snp' | 0.004 | synonymous_variant c.801G>A p.Leu267Leu | |
| '3322398','G:A','snp' | 0.004 | synonymous_variant c.825G>A p.Leu275Leu | |
| '3405147','A:T','snp' | 0.004 | | |

#### Klebsiella pneumoniae - tobramycin

| Bacteria | Antibiotic | F1 Score | MCC |
|---|---|---|---|
| Klebsiella pneumoniae | tobramycin | 0.893 | 0.778 |

Table 70: Feature Importance for Klebsiella pneumoniae - tobramycin

| Feature Name | Gini Importance | Mutation Annotation | Gene |
|---|---|---|---|
| '2963641','G:A','snp' | 0.007 | synonymous_variant c.585G>A p.Ala195Ala | |
| L5EHL7HU_CDMPAKHP_05726_Aminoglycoside_N_6___acetyltransferase_type_1 | 0.004 | | |
| '3065847','A:G','snp' | 0.004 | | |
| '2412656','T:C','snp' | 0.004 | synonymous_variant c.286T>C p.Leu96Leu | |
| 05GCBKNR_FPLHKBIC_00303_hypothetical_protein | 0.004 | | |

**Table 70 – continued from previous page**

| Feature Name | Gini Importance | Mutation Annotation | Gene |
|---|---|---|---|
| 001QJFFS_HPLJBCIE_00039_tetracycline_repressor_protein_class_G | 0.004 | | |
| ’2882298’,’C:T’,’snp’ | 0.004 | synonymous_variant c.1002G>A p.Gln334Gln | |
| ’5260577’,’A:G’,’snp’ | 0.004 | synonymous_variant c.130T>C p.Leu44Leu | |
| ’3286798’,’G:A’,’snp’ | 0.004 | | |
| ’2603975’,’C:T’,’snp’ | 0.004 | missense_variant c.841C>T p.Pro281Ser | |

#### Salmonella enterica - chloramphenicol

| Bacteria | Antibiotic | F1 Score | MCC |
|---|---|---|---|
| Salmonella enterica | chloramphenicol | 0.898 | 0.874 |

Table 71: Feature Importance for Salmonella enterica - chloramphenicol

| Feature Name | Gini Importance | Mutation Annotation | Gene |
|---|---|---|---|
| 00Q9M4HG_BPCMGPCK_01806_Inner_membrane_transport_protein_YdhC | 0.021 | | |
| 00WQ11Z9_IBABPKKH_02379_hypothetical_protein | 0.009 | | |
| 00Q9M4HG_BPCMGPCK_02897_hypothetical_protein | 0.009 | | |
| 00WQ11Z9_IBABPKKH_04718_putative_multidrug_efflux_protein | 0.009 | | |
| 00WQ11Z9_IBABPKKH_04294_IS6_family_transposase_IS6100 | 0.009 | | |
| 00ZGQ2GU_NJPCPHLH_04907_60_kDa_chaperonin | 0.009 | | |
| 00ZGQ2GU_NJPCPHLH_04400_Tyrosine_recombinase_XerC | 0.008 | | |
| 00WQ11Z9_IBABPKKH_02406_hypothetical_protein | 0.008 | | |

**Table 71 – continued from previous page**

| Feature Name | Gini Importance | Mutation Annotation | Gene |
|---|---|---|---|
| 19DPBHKE_HAPMAMJP_03441_hypothetical_protein | 0.008 | | |
| '2417680','G:T','snp' | 0.008 | missense_variant c.997C>A p.Leu333Ile | menF |

#### Salmonella enterica - gentamicin

| Bacteria | Antibiotic | F1 Score | MCC |
|---|---|---|---|
| Salmonella enterica | gentamicin | 0.759 | 0.734 |

Table 72: Feature Importance for Salmonella enterica - gentamicin

| Feature Name | Gini Importance | Mutation Annotation | Gene |
|---|---|---|---|
| 00ZGQ2GU_NJPCPHLH_04400_Tyrosine_recombinase_XerC | 0.013 | | |
| 7IRHR44U_EIMMAFOK_03916_hypothetical_protein | 0.011 | | |
| MUD4U9AA_KOMBHBOF_01768_hypothetical_protein | 0.009 | | |
| K1EJ2S7Q_NMEGFFKJ_04055_hypothetical_protein | 0.008 | | |
| 72MFX2FO_JOMKBDPJ_04903_hypothetical_protein | 0.007 | | |
| 00WQ11Z9_IBABPKKH_04770_putative_membrane_transporter_of_cations | 0.006 | | |
| 2NQ2WNLA_EBIBFBDP_04091_Protein_PndA | 0.006 | | |
| YVHVP2M1_LJECKDLM_04639_hypothetical_protein | 0.006 | | |
| 0BCHF2R4_OCHDPHLH_02836_hypothetical_protein | 0.005 | | |
| 03BKKDIO_MMMFKGBG_04455_hypothetical_protein | 0.005 | | |

#### Salmonella enterica - ceftiofur

| Bacteria | Antibiotic | F1 Score | MCC |
|---|---|---|---|
| Salmonella enterica | ceftiofur | 0.930 | 0.913 |

Table 73: Feature Importance for Salmonella enterica - ceftiofur

| Feature Name | Gini Importance | Mutation Annotation | Gene |
|---|---|---|---|
| 00Q9M4HG_BPCMGPCK_03608_outer_membrane_lipoprotein | 0.018 | | |
| 00Q9M4HG_BPCMGPCK_03609_penicillin__binding_protein | 0.014 | | |
| 00Q9M4HG_BPCMGPCK_03610_hypothetical_protein | 0.011 | | |
| RH7QGROV_BNEGCPGP_04477_IS1380_family_transposase_ISEcp1 | 0.008 | | |
| 00Q9M4HG_BPCMGPCK_03607_putative_DMT_superfamily_transport_protein | 0.007 | | |
| 03QPFX4M_JGBABALF_04112_hypothetical_protein | 0.007 | | |
| 00Q9M4HG_BPCMGPCK_02942_hypothetical_protein | 0.006 | | |
| 00Q9M4HG_BPCMGPCK_02919_hypothetical_protein | 0.006 | | |
| 00Q9M4HG_BPCMGPCK_02953_hypothetical_protein | 0.006 | | |
| 03BKKDIO_MMMFKGBG_04452_hypothetical_protein | 0.005 | | |

#### Salmonella enterica - sulfisoxazole

| Bacteria | Antibiotic | F1 Score | MCC |
|---|---|---|---|
| Salmonella enterica | sulfisoxazole | 0.891 | 0.841 |

Table 74: Feature Importance for Salmonella enterica - sulfisoxazole

| Feature Name | Gini Importance | Mutation Annotation | Gene |
|---|---|---|---|
| 044SS4JL_FCCMCGPF_03841_hypothetical_protein | 0.011 | | |
| '2217995','A:G','snp' | 0.011 | missense_variant c.2473A>G p.Ser825Gly | yegN |
| 00Q9M4HG_BPCMGPCK_02950_hypothetical_protein | 0.009 | | |
| '3586880','G:T','snp' | 0.008 | synonymous_variant c.486C>A p.Val162Val | secY |
| '3449458','C:A','snp' | 0.007 | synonymous_variant c.1932G>T p.Gly644Gly | pnp |
| 00Q9M4HG_BPCMGPCK_02943_DNA_binding_protein_HU_beta__NS1__HU_1 | 0.006 | | |
| '227908','A:T','snp' | 0.006 | missense_variant c.248A>T p.Gln83Leu | fhuB |
| 08NBTB2V_PBFNHNIL_04032_IS256_family_transposase_ISEc58 | 0.006 | | |
| 00ZGQ2GU_NJPCPHLH_04400_Tyrosine_recombinase_XerC | 0.006 | | |
| 00Q9M4HG_BPCMGPCK_02918_hypothetical_protein | 0.005 | | |

#### Salmonella enterica - streptomycin

| Bacteria | Antibiotic | F1 Score | MCC |
|---|---|---|---|
| Salmonella enterica | streptomycin | 0.900 | 0.827 |

Table 75: Feature Importance for Salmonella enterica - streptomycin

| Feature Name | Gini Importance | Mutation Annotation | Gene |
|---|---|---|---|
| 00Q9M4HG_BPCMGPCK_03416_Aminoglycoside_3__phosphotransferase | 0.010 | | |
| 00WQ11Z9_IBABPKKH_04295_hypothetical_protein | 0.005 | | |
| 00WQ11Z9_IBABPKKH_04215_hypothetical_protein | 0.005 | | |

**Table 75 – continued from previous page**

| Feature Name | Gini Importance | Mutation Annotation | Gene |
|---|---|---|---|
| 00ZGQ2GU_NJPCPHLH_04400_Tyrosine_recombinase_XerC | 0.005 | | |
| 00WQ11Z9_IBABPKKH_04213_hypothetical_protein | 0.004 | | |
| RNMOCWEB_ABCGGBDN_04742_hypothetical_protein | 0.004 | | |
| ZH9AGC7H_KNCCIBNF_04381_hypothetical_protein | 0.004 | | |
| '2048377','C:T','snp' | 0.004 | missense_variant c.766G>A p.Glu256Lys | fliC |
| 00WQ11Z9_IBABPKKH_02379_hypothetical_protein | 0.003 | | |
| '1174580','T:A','snp' | 0.003 | synonymous_variant c.336A>T p.Gly112Gly | yccA |

#### Salmonella enterica - ampicillin

| Bacteria | Antibiotic | F1 Score | MCC |
|---|---|---|---|
| Salmonella enterica | ampicillin | 0.887 | 0.806 |

Table 76: Feature Importance for Salmonella enterica - ampicillin

| Feature Name | Gini Importance | Mutation Annotation | Gene |
|---|---|---|---|
| '1804950','A:C','snp' | 0.006 | synonymous_variant c.2475T>G p.Leu825Leu | acnA |
| '94669','C:T','snp' | 0.005 | | |
| RH7QGROV_BNEGCPGP_04477_IS1380_family_transposase_ISEcp1 | 0.005 | | |
| '4268203','G:C','snp' | 0.005 | synonymous_variant c.261G>C p.Gly87Gly | |
| 00ZGQ2GU_NJPCPHLH_04956_putative_resolvase | 0.005 | | |
| 00Q9M4HG_BPCMGPCK_03607_putative_DMT_superfamily_transport_protein | 0.005 | | |
| 00ZGQ2GU_NJPCPHLH_04957_Beta_lactamase_TEM | 0.004 | | |

**Table 76 – continued from previous page**

| Feature Name | Gini Importance | Mutation Annotation | Gene |
|---|---|---|---|
| 00Q9M4HG_BPCMGPCK_03608_outer_membrane_lipoprotein | 0.004 | | |
| '3034095','T:C','snp' | 0.003 | synonymous_variant c.258A>G p.Ser86Ser | spaP |
| '2234484','A:G','snp' | 0.003 | synonymous_variant c.474A>G p.Ala158Ala | yegS |

#### Salmonella enterica - cefoxitin

| Bacteria | Antibiotic | F1 Score | MCC |
|---|---|---|---|
| Salmonella enterica | cefoxitin | 0.870 | 0.847 |

Table 77: Feature Importance for Salmonella enterica - cefoxitin

| Feature Name | Gini Importance | Mutation Annotation | Gene |
|---|---|---|---|
| 00Q9M4HG_BPCMGPCK_03607_putative_DMT_superfamily_transport_protein | 0.023 | | |
| RH7QGROV_BNEGCPGP_04477_IS1380_family_transposase_ISEcp1 | 0.015 | | |
| 00Q9M4HG_BPCMGPCK_03608_outer_membrane_lipoprotein | 0.014 | | |
| 00Q9M4HG_BPCMGPCK_03610_hypothetical_protein | 0.013 | | |
| 00Q9M4HG_BPCMGPCK_03609_penicillin__binding_protein | 0.009 | | |
| 00Q9M4HG_BPCMGPCK_02926_hypothetical_protein | 0.009 | | |
| 00Q9M4HG_BPCMGPCK_02949_hypothetical_protein | 0.009 | | |
| 00Q9M4HG_BPCMGPCK_02948_hypothetical_protein | 0.008 | | |
| 00Q9M4HG_BPCMGPCK_02942_hypothetical_protein | 0.008 | | |
| 0HMOA7IN_OADOOPHH_04745_hypothetical_protein | 0.008 | | |

#### Salmonella enterica - ceftriaxone

| Bacteria | Antibiotic | F1 Score | MCC |
|---|---|---|---|
| Salmonella enterica | ceftriaxone | 0.906 | 0.883 |

Table 78: Feature Importance for Salmonella enterica - ceftriaxone

| Feature Name | Gini Importance | Mutation Annotation | Gene |
|---|---|---|---|
| 00Q9M4HG_BPCMGPCK_03607_putative_DMT_superfamily_transport_protein | 0.022 | | |
| 00Q9M4HG_BPCMGPCK_02923_hypothetical_protein | 0.012 | | |
| 00Q9M4HG_BPCMGPCK_03609_penicillin__binding_protein | 0.010 | | |
| 03QPFX4M_JGBABALF_04112_hypothetical_protein | 0.010 | | |
| 00Q9M4HG_BPCMGPCK_02936_hypothetical_protein | 0.009 | | |
| 00Q9M4HG_BPCMGPCK_02922_hypothetical_protein | 0.009 | | |
| RH7QGROV_BNEGCPGP_04477_IS1380_family_transposase_ISEcp1 | 0.008 | | |
| 00Q9M4HG_BPCMGPCK_03429_hypothetical_protein | 0.008 | | |
| 00Q9M4HG_BPCMGPCK_02912_hypothetical_protein | 0.008 | | |
| 0P3QTX4R_MEMEGJBF_04152_hypothetical_protein | 0.007 | | |

#### Salmonella enterica - tetracycline

| Bacteria | Antibiotic | F1 Score | MCC |
|---|---|---|---|
| Salmonella enterica | tetracycline | 0.925 | 0.838 |

Table 79: Feature Importance for Salmonella enterica - tetracycline

| Feature Name | Gini Importance | Mutation Annotation | Gene |
|---|---|---|---|
| 00Q9M4HG_BPCMGPCK_02954_IS6_family_transposase_IS26 | 0.008 | | |
| 249RYVAR_NKDGCMPF_03630_hypothetical_protein | 0.007 | | |
| 00Q9M4HG_BPCMGPCK_03412_putative_multidrug_efflux_protein | 0.006 | | |
| '3793981','A:G','snp' | 0.005 | synonymous_variant c.3468T>C p.Val1156Val | yhjL |
| '253485','G:C','snp' | 0.005 | synonymous_variant c.363C>G p.Leu121Leu | map |
| '2099781','A:C','snp' | 0.005 | missense_variant c.1419T>G p.Asp473Glu | cbiP |
| '758353','T:C','snp' | 0.005 | synonymous_variant c.657A>G p.Ala219Ala | ybfF |
| '1424956','C:A','snp' | 0.005 | missense_variant c.1393C>A p.Arg465Ser | ydiU |
| '4530315','A:G','snp' | 0.005 | synonymous_variant c.204T>C p.Gly68Gly | phnA |
| '564240','G:C','snp' | 0.004 | missense_variant c.633C>G p.Asp211Glu | ybbN |
